# Annotation Comparison Explorer (ACE): connecting brain cell types across studies of health and Alzheimer’s Disease

**DOI:** 10.1101/2025.02.11.637559

**Authors:** Jeremy A. Miller, Kyle J. Travaglini, Tain Luquez, Rachel E. Hostetler, Aaron Oster, Scott Daniel, Bosiljka Tasic, Vilas Menon

## Abstract

**Background:** Single-cell multiomic technologies have allowed unprecedented access to gene profiles of individual cells across species and organ systems, including the brain. The Allen Institute has created foundational atlases characterizing mammalian cell types in the adult mouse brain and the neocortex of humans with and without Alzheimer’s disease (AD). However, proliferation of public cell type classifications (or ‘taxonomies’) by us and others creates a challenge for knowledge integration.

**Results:** Here, we introduce Annotation Comparison Explorer (ACE), a web application for comparing cell type assignments and other cell-based annotations (e.g., donor demographics, anatomic locations, quality control metrics). ACE can filter cells and includes an interactive set of visualization tools, plot and data downloads, and statistics for comparing two or more taxonomy annotations alongside collected knowledge (e.g., cell type aliases, marker genes, abundance changes in disease). In this study we describe ACE functionality and present the following three ACE use cases. First, we demonstrate how a user can assign labels from the Seattle Alzheimer’s Disease Brain Cell Atlas (SEA-AD) taxonomy to their own cells and compare these ‘mappings’ to user-defined cell type assignments and other cell metadata, using a previous cell type classification from the Allen Institute and two independent studies of different brain diseases as inputs. Second, we extend this approach for comparison of ten published human AD studies previously reprocessed through a common data analysis pipeline, and identify congruent cell type abundance changes in AD, including a decrease in certain somatostatin interneurons. Finally, ACE includes translation tables between different mouse and human brain cell type taxonomies on Allen Brain Map, from initial studies in neocortex to more recent studies spanning the whole brain, along with a human immune cell atlas focused on peripheral blood mononuclear cells. These use cases represent three of many possible applications for ACE.

**Conclusions:** ACE combines standard and custom visualizations into a user-friendly, open-source web tool for exploring categorical and numeric relationships and translating cell type classifications and knowledge across studies. ACE can be freely and publicly accessed at https://sea-ad.shinyapps.io/ACEapp/.

## Background

Since their invention more than a decade ago, single cell [1] and single nucleus [2] RNA-sequencing technologies (sc/snRNA-seq) have revolutionized the definition of cell types from a largely qualitative to a highly quantitative concept. Large scale, community efforts from the Human Cell Atlas [3] and others now aim to comprehensively define and characterize all cell types in the human body. In the brain, cell types historically defined by their shape, firing properties, or expression of at most a few marker genes, can now be defined by the molecular content of their somas or nuclei. The Allen Institute has created foundational atlases characterizing brain cell types in the adult mammals [4–11], with more than 3000 cell types in human [11] and more than 5000 cell types in mouse [4] defined to date, finding that molecularly-defined types from sc/snRNA-seq largely (but not entirely) align with historically-defined types and tend to be anatomically constrained. While most studies still define their own cell type classifications (or “taxonomies”), large-scale efforts to define integrated reference atlases in brain [4–11] and other organ systems (e.g., lung [12]), provide comprehensive transcriptomic cell type definitions in the context of extensive multi-modal knowledge. In human and mouse neocortex, for example, morphoelectric properties of cells can be inferred by aligning Patch-seq studies [13–18] with existing cell type taxonomies, and spatial locations of all mouse brain cell types have been established using spatial transcriptomics [4]. These cell type taxonomies have proven to be highly conserved across data collected on different platforms, across multiple data modalities, and even between species where few homologous marker genes show conserved patterns [6, 7, 10, 19]. Sc/snRNA-seq can also identify changes in cell type abundance, cell state, or gene expression profiles in aging [20] and disease, including Alzheimer’s Disease (AD) [21–31], where several groups have identified novel microglial and other glial states associated with AD pathology [28, 31–33]. Collectively, these efforts present brain cells as a complex landscape of intersecting molecular cell types and cell states, which neuroscientists are slowly coming to understand.

With so many public cell type classifications available and many groups choosing to define their own, linking cell types and associated knowledge between studies remains a major challenge. Many studies define their own classifications using inconsistent nomenclature. For example, three recent studies that identified microglial types with increased abundance in AD defined them as Mic. 12/13 [31], MG1 [28], and Micro-PVM_3-SEAAD [21] using different abbreviations for microglia and a different system for numbering cell types. Furthermore, cell typing studies often have different levels of resolution depending on the number of cells collected, the quality of the data, or the focus of the study. For example, these same microglial studies arrived at very different numbers of total microglial populations: 16, three, and six, respectively. Furthermore, many studies of disease break neurons into a handful of broad groups, while atlases in the healthy brain identify several dozen neuronal types per region. Public tools like MapMyCells (RRID:SCR_024672) and Azimuth [34] provide user-friendly interfaces for scientists to align data to reference taxonomies, and recent efforts have attempted to standardize nomenclature in brain cell types [35] for research groups committed to defining their own cell types. Even as these efforts gain traction, linking information from newer, larger studies to highly cited and well-annotated studies on focused brain regions and cell types remains a challenge.

To address these challenges we have developed Annotation Comparison Explorer (ACE; RRID:SCR_026496), a web and associated R shiny application for comparison of two or more annotations such as (i) cell type assignments (e.g., from different mapping/clustering algorithms), (ii) donor metadata (e.g., donor, sex, age), and (iii) cell metadata (e.g., anatomic location, quality control (QC) metrics). ACE comes prepopulated with several highly requested translations between published brain cell types in health and disease, and also allows users to provide their own tables for comparing annotations. Here we describe ACE functionality and present common use cases. ACE is freely and publicly available for use as a web application (https://sea-ad.shinyapps.io/ACEapp/) and on GitHub (github.com/AllenInstitute/ACE).

### Implementation

#### Overview of ACE

ACE compares cell type annotations across datasets in three sequential steps. First, a data set must be selected, either by uploading user data or by choosing a pre-defined table of public cell type data. Second, the data set can optionally be filtered to focus on specific annotations.

Finally, filtered data can be explored through a variety of interactive visualizations and statistics. ACE also contains an informational pane with links to additional resources for help or to contribute, including an informational webinar, a video introduction, video use cases, and an extensive user guide (**Supplemental Text 1**). As a simple example (**Supplemental Figure 1**), eight pyramidal neurons are initially classified into two groups based on size. Addition of a ninth neuron leads to a reclassification into three groups, resulting in two different size annotations for the original eight neurons. After saving the data in a simple format and uploading to ACE, ACE can show how size classifications change between the two annotations using river plots, revealing shifts in the definitions of size categories.

#### Creation and formatting of data annotation tables

As of October 2025, ACE includes 18 pre-built tables largely focused on mammalian brain cell types (see **Supplemental Table 1**), although the tool can be generalized to a wide range of additional applications. These tables include comparisons for disease studies spanning six brain diseases and 22 independent data sets [21–31, 36], cell type classifications from human [7, 9–11, 21] and mouse [4, 5, 8, 19, 20] brain and from human blood [37], and electrophysiological properties of human [16–18] and mouse [14, 15] neocortical cells collected using Patch-seq (a technique that links these properties to gene-based cell types [38]). All pre-built tables are available on GitHub and include an informative project description (**Supplemental Table 1**). To the extent possible, these tables derive from publicly available sources, with the exception that several of the files from AD studies come from controlled access materials or were provided directly from manuscript authors and therefore cannot be publicly shared. The ACE GitHub repository also includes a “data_ingest_scripts” folder containing scripts for converting these source files to the data tables used in ACE (**Supplemental Table 1**). For publicly-accessible sources, direct URLs to files are included in the data ingest scripts, while for non-public files the means of access is indicated.

For both user-provided and pre-built data sets, ACE runs off two input tables: a required data (e.g., cell) table and an optional annotation (e.g., cell type) table. The data table encodes data points (e.g., cells) as rows and various cell annotations (e.g., cell type assignments, anatomic structures, QC metrics) as columns, and is the table from which nearly all of the visualizations and statistics derive. This can be provided either as a standard text form (‘.csv’ or ‘.csv.gz’) or in common formats for accessing snRNA-seq data in R and python (‘.h5ad’, in which case only the ‘obs’ slot is read, or ‘.feather’). ACE can take any cell-level information (either numeric or categorical) as input; however, it is worth noting that gene expression values are typically not provided to ACE (see **Discussion**). The annotation table encodes additional information about each annotation and, if provided, must be a ‘.csv’ or ‘.csv.gz’ file where rows represent individual metadata values (e.g., cell type names) and columns are any information about those metadata values. Two columns have special meaning: ‘cell_type’ holds the names of the metadata items in the data table and dictates the order of annotations in ACE visualizations, and ‘direction’ indicates changes in abundance with disease (‘Up with disease’, ‘Down with disease’, ‘No change’, ‘Not assessed’ or ‘Not provided’). All other information will be presented to a user when the relevant annotation is selected in the ‘Explore individual annotations’ tab, and there are no other specific restrictions on what goes in the rows or columns of this table.

#### Components of the ACE application

ACE is divided into three primary panels, mirroring the steps introduced above. The “Upload or select data set” panel allows users to upload their own data or choose from one of the pre-built annotation comparison tables described above (**Figure 1A**). To explore user-provided data, either click on the “Browse…” buttons to upload a file or enter the URL of a file in a public source (e.g., on GitHub). Pre-built files can be accessed by clicking first on the dropdown box to “Choose a category” and then selecting an existing data set from the other drop-down box.

**Figure 1:**
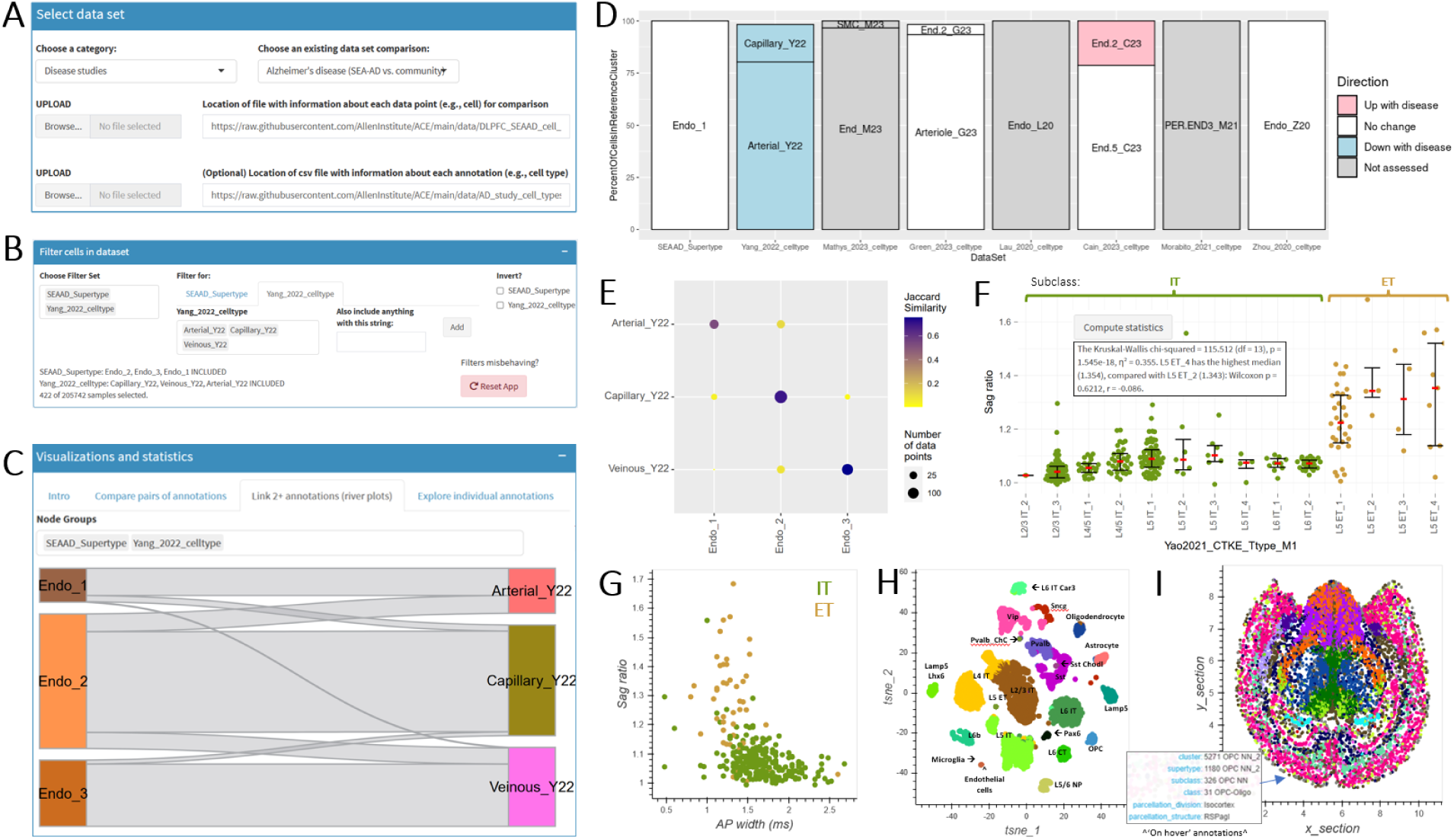
Overview of ACE functionality. **A-C)** Screenshots of the three main components of ACE as applied to comparison of endothelial cell types between two studies of AD. The top panel (**A**) allows a user to select from a predefined data set (as shown here) or upload their own. Description text omitted for space. The middle panel (**B**) allows a user to filter data points within the selected data set based on one or more annotation columns, in this case by only including cells labeled as endothelial types in the SEAAD and Yang_2022 studies. The bottom panel (**C**) includes tabs for all visualizations and analysis performed by ACE on the filtered dataset, in this case showing the relationship between annotations of three endothelial cell types in SEAAD and Yang_2022. **D-I)** Visualizations available in ACE. **D**) Custom bar plots showing the relationship between single value (Endo_1) in one annotation (SEAAD_Supertype) and values in one or more other annotations (cell type assignments from other studies), and color-coding by whether these values show a change in abundance with AD. **E**) Confusion matrices showing the Jaccard similarity between all values in a pair of annotations, in this case showing the same information as in **C**. **F**) Bee swarm plots showing any numeric property (Sag ratio) divided by values of one annotation (Yao2021_CTKE_Type_M1) and color-coded by another (Subclass). Inset shows output when the “Compute statistics” button is pressed in this view. **G-I**) Scatterplots showing the relationship between two numeric annotations such as electrophysiological parameters (**G**), UMAP coordinates (**H**), or physical cell locations (**I**), and color-coding by a categorical annotation (Subclass). Subclass assignments in **G** and **H** are manually added and the inset in **I** shows ‘on hover’ annotations for the indicated cell.

The “Filter cells in dataset” panel allows users to restrict the input data to specific annotations (e.g., endothelial cells rather than all cells) (**Figure 1B**). The “Choose Filter Set” box lets users select one or more metadata columns for filtering, with autocomplete functionality to aid in selection. For each selected categorical (text) metadata column, the “Filter for” box allows filtering to include only specific values (with an option to invert the selection) or include specific text strings. For each selected numeric metadata column, a user can filter by setting a range, and a histogram of values is shown above to aid in filtering. Details about the current selection are also shown in the app.

The “Visualization and statistics” panel provides a variety of interactive options for exploration of the relationships between selected data points, which are described in the default “Intro” tab. For example, river plots show relationships between two or more categories arranged in columns. The “rivers” connecting adjacent columns show the number of items (e.g., cells) shared between categories, with thicker rivers representing larger overlaps. After filtering for endothelial cells (**Figure 1B**), a clear relationship between supertypes from the Seattle Alzheimer’s Brain Cell Atlas (SEA-AD; RRID:SCR_023110) [21] and reported arterial, capillary, and venous vascular cell types [29] can be seen (**Figure 1C**), suggesting obvious improvements to future cell type annotations. ACE also provides novel functionality for focused exploration of individual annotations; for example, arterial cells represent a distinct cell population across multiple studies, with the great majority of all cells labeled as “Endo_1” also being labeled to a single cell type in seven other AD studies (**Figure 1D**). A confusion matrix can be used to show the correspondence between two different sets of classifications (in **Figure 1E**, cell type assignments from two different studies). The matrix displays how many items (cells) assigned to a particular category in one classification are assigned to each category in the other classification, and the size and color of points can be used to encode different types of information like Jaccard similarity, fraction representation, or cell counts. When comparing two annotations, confusion matrices and river plots show the same information in different ways.

ACE can also visualize numeric variables (**Figure 1F-I**). For example, electrophysiological differences between intratelencephalic (IT) and extratelencephalic (ET) cells are well established with ET neurons having a higher ratio of voltage at the peak and voltage at steady-state in response to hyperpolarizing current injections (“Sag ratio”) than IT neurons in humans [13]. By exploring an available table of Patch-seq data collected in mouse primary motor cortex [14] and restricting the view to these two subclasses (ET and IT), this difference is clearly consistent between all ET and IT cell types (**Figure 1F**), shows a statistical difference across cell types (**Figure 1F**; inset), and matches previous reports in humans [13]. Additional separation between mouse ET and IT cells can be seen when comparing Sag ratio (higher in ET) and action potential (AP) width (defined as the duration of the AP at half-amplitude from AP threshold; higher in IT) using a scatter plot (**Figure 1G**). While not its primary use, ACE’s “Compare numeric annotations” can also be used to overlay annotations on latent representations of sc/snRNA-seq data (**Figure 1H**; in human middle temporal gyrus, MTG) or on physical cell locations in tissue slices (**Figure 1I**; in a mouse coronal section). Mouse cell types show clear spatial patterning in the brain, as previously reported [4, 39].

#### Creation of a hosted R shiny application

ACE was created using R shiny (RRID:SCR_001626) [40], an extension of the R programming language supporting interactive analysis and visualization, and then uploaded to shinyapps.io so it is accessible anywhere on the web without the need for users to install anything on their computer or use R. For users preferring a local instance of the tool, ACE can also be installed using rig [41] for R installation management along with the R package renv [42] to manage project-specific libraries and dependencies, ensuring reproducibility between shinyapps.io and local environments (see instructions on GitHub). As detailed above, ACE can read annotation tables from h5ad, feather, and most commonly text files (csv and csv.gz) which can be quickly uploaded using vroom [43]. Visualizations were created using a combination of custom functions and functions available as part of the ggplot2 (RRID:SCR_014601) [44] and the archived rbokeh [45] R package (v0.5.2), which we host at https://github.com/jeremymiller/rbokeh for reproducibility. Web app usage is being tracked using Google Analytics; no user information (beyond city of origin) or uploaded data is retained for any purpose. A video webinar overviewing ACE, a set of short videos introducing ACE and describing specific use cases, and a detailed written user guide have been created to improve web application usability and are included alongside link outs to related tools and ways to contribute on the left menu panel.

#### Generating annotation tables for cell type classification in healthy adult brain

ACE includes several predefined annotation tables comparing cell type classifications across public studies of mouse and human brain, including neocortical areas, hippocampus, basal ganglia, and whole brain. Each table (or “dataset”) includes a description describing the specific taxonomies and annotations included therein. All of these tables except the ones noted below were generated by (1) identifying a set of studies that were performed using an overlapping set of cells, (2) subsetting existing cell annotation tables from each study to only include these common cells, and (3) compiling relevant cell annotations (e.g., cell type assignments, brain region, and donor information) from each study into a single table. Large data sets were subsampled to less than ∼500,000 cells to speed up visualizations. Numeric values such as UMAP coordinates, physical spatial locations, and mapping confidence scores were also included as relevant. The “Middle temporal gyrus (initial studies)” table connected taxonomies from two distinct studies of MTG by using MapMyCells (RRID:SCR_024672; https://portal.brain-map.org/atlases-and-data/bkp/mapmycells) to transfer labels from the recent human MTG SEA-AD taxonomy [21] to cells in an earlier MTG taxonomy [7] that has been used as a reference for several human Patch-seq studies using deep generative mapping. This process is discussed in detail below. Similarly, the “Mammalian Basal Ganglia Consensus Cell Type Atlas” table includes cell annotations along with label transfer from both whole human brain and whole mouse brain using the default hierarchical mapping approach.

#### Generating annotation tables comparing cell types across studies of Alzheimer’s disease

To link annotations across 11 studies of AD, we integrated data from ten community-based studies of AD focused on dorsolateral prefrontal cortex (DFC) [22–31] with DFC data from SEA-AD [21] and then compared transferred SEA-AD labels with published information for each study (**Supplemental Figure 2A**). Integration and label transfer was performed as described in [21]. Briefly, raw sequencing data was downloaded from the AD Knowledge Portal hosted on Synapse (RRID:SCR_006307) or from the Sequencing Read Archive (SRA; RRID:SCR_004891) and processed using the same pipeline as was used for SEA-AD snRNA-seq data. Processed data were similarly QCed and then mapped to SEA-AD cell types using a variation of the Deep Generative Mapping model on MapMyCells, which is only available for this cell type taxonomy (Hierarchical Mapping is the default mapping algorithm for all other taxonomies). Cell type classifications from the original studies were collected from Synapse, original manuscripts, and email correspondence with authors and linked to mapped cell type using unique cell identifiers, only retaining cells with both SEA-AD and initial cell type assignments for additional processing.

ACE assumes each data point represents a single cell with multiple annotations to the same cells. Since these AD studies include pairs of annotations for many different sets of cells, we applied a novel approach to link cluster assignments from community data sets to SEA-AD cells (**Supplemental Figure 2B**). For each community study, we first generated a confusion matrix comparing the reported cluster assignments and the mapped SEA-AD cell types (supertypes). We then assigned each SEA-AD cell a cell type from the external study based on the likelihood of cells from the SEA-AD type being assigned to a given cell type in the external study. SEA-AD cells with supertypes that were not found in a given study were assigned a cell type of “noMappedCells” (e.g., when comparing neuronal cells to studies focused on glial cell types). This process was then repeated for each of the 10 studies. Cell type assignments for broader cell type definitions were inferred from their higher-resolution counterparts. For example, cells assigned to “Sst 25” were inferred to be “Sst” cells and “GABAergic interneurons”.

Cell types from all 11 studies were also assigned a direction change in abundance with AD based exclusively on reported findings from initial publications (**Supplemental Figure 2B**). Specifically, we reviewed both the text and figures from all relevant manuscripts [21–31] for quantitative values (when available) or for qualitative statements (when values were not presented) regarding a cell type’s change in abundance with AD and then recorded this information in the cell type annotation table along with notes indicating how direction changes were defined. Cell types where abundance was assessed were listed as “Down with disease,” “Up with disease,” or “No change”. We also indicated cases where this information was “Not assessed” or “Not provided.”

#### Generating annotation tables for diverse data sets

While ACE was initially created for linking annotations between cell type taxonomies on Allen Brain Map and for comparing studies of AD, the tool is highly versatile and has since been extended for diverse purposes. First, annotation tables from five Patch-seq studies are included [14–18]. These studies all include cell type classifications defined in the original publication, typically using label transfer from existing mouse or human neocortical cell type taxonomies along with morphoelectric characteristics and other cell or donor metadata measured for each cell. With this information ACE can be used to visualize and statistically assess differences in cellular characteristics between neuronal cell types and how various morphoelectric features relate. Second, we have included additional studies of aging and disease: a study of aged mice [20] where cells are assigned common types from adult whole brain allows direct comparison of cell type abundances with age, while two additional studies in human disease map cell types from human DFC and whole brain allow direct comparison between AD, Parkison’s disease (PD), autism spectrum disorder (ASD), major depressive disorder (MDD), post-traumatic stress disorder (PTSD), and William’s syndrome [36]. Finally, we include a study from the Allen Institute for Immunology focused on cell type classification in peripheral blood mononuclear cells (PBMCs) [37], demonstrating the utility of ACE on applications outside of the brain.

## Results

In the following sections we present several use cases for ACE, which all rely on the pairs of simple cell and cell type annotation tables detailed above. One use case details how to integrate clustering and mapping results for user-defined data, while the rest rely on pre-built data tables provided in ACE or as supplemental materials.

### Use Case 1: Comparing user-provided clustering and MapMyCells mapping results

The Allen Institute has developed several resources for cross-study analysis and label transfer to facilitate the transfer of this knowledge to other groups, including: (1) multiple high-quality and well-annotated cell type taxonomies [4–11, 19, 21] stored in standard formats and with public data visualization tools (e.g., the Allen Brain Cell Atlas (RRID:SCR_024440)); (2) interactive and code-based algorithms for assigning cell types from these taxonomies to user data (e.g., MapMyCells), and (3) tools for visualization and exploration of cell type assignments across studies (e.g., ACE). In this use case we demonstrate how to identify reference brain cell types associated with user-provided sc/snRNA-seq data and compare these results to user metadata without the need to code (**Figure 2A**). This process involves the following steps. First, collect a cell by gene matrix in a standard format (h5ad or csv) and a cell annotation table that includes pre-defined cluster calls. Next, run MapMyCells to transfer cell types from a user-provided taxonomy of interest using one of three available algorithms. Detailed instructions for running MapMyCells are available on Allen Brain Map, but this involves uploading a cell by gene matrix, clicking a few options in a graphical user interface (GUI), and then downloading and unzipping results. Third, join the mapping results downloaded from MapMyCells with the initial cell annotation table to create a single file containing all cell annotations, for example by copying columns between files in Excel. Finally, upload this joined file (along with any relevant cell type annotations as an optional second csv file) into ACE and explore.

**Figure 2:**
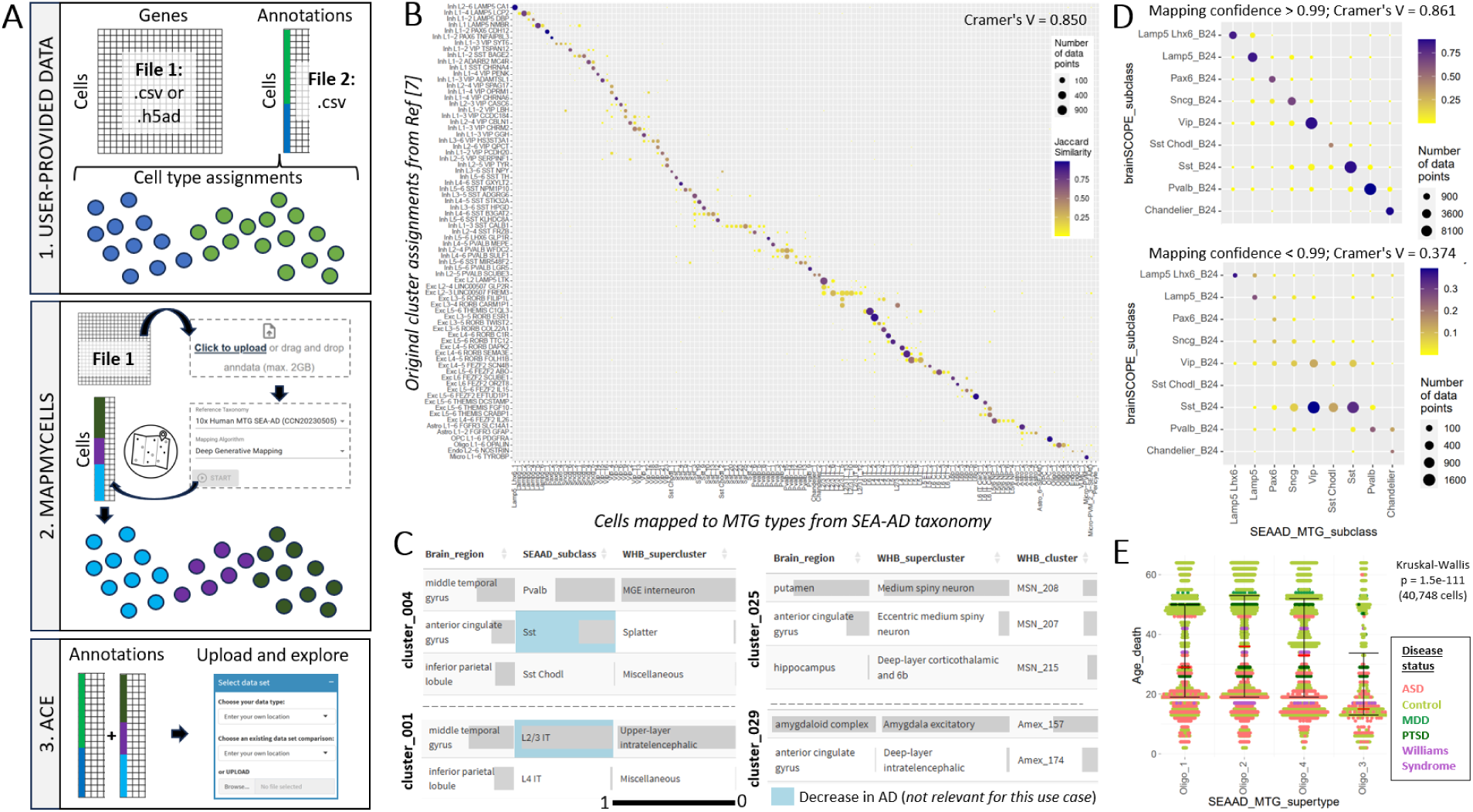
Comparison of user and reference brain cell types using MapMyCells and ACE. **A**) Overview of user workflow for a code-free comparison of user-provided gene expression data and associated cell annotations with cell type mapping results using MapMyCells. **B**) Confusion matrix after using this workflow to map gene expression data from Hodge, Bakken et al 2019 [7] to the SEA-AD taxonomy. These results are provided as a predefined table in ACE (“Middle temporal gyrus (initial studies)”), with the workflow for mapping Hodge to SEA-AD available on Allen Brain Map at https://portal.brain-map.org/atlases-and-data/bkp/mapmycells/mapmycells-use-case-single-nucleus-rnaseq-from-human-mtg and a detailed conversion script available on GitHub (**Supplemental Table 1**). **C**) Table bar graph representations for four PD clusters (indicated on y axis label) showing the fraction of cells in that cluster collected from different brain region dissections (left columns) or mapping to different cell types (center and right columns). In all cases, grey bars represent fractions of cells of a given column, with columns summing to 1. **D**) Confusion matrix comparing brainSCOPE clustering (y-axis) and SEA-AD mapping (x-axis) results for GABAergic interneurons in DFC for cells with high (top) vs. lower (bottom) confidence mapping. **E**) Beeswarm with overlaid box and whisker plot showing a decrease in expression in cells mapping to Oligo_3 relative to other oligodendrocyte supertypes with age. Cells color-coded by disease status do not show an obvious difference in age effect.

To demonstrate the utility of this workflow, we took data from an early classification of MTG that has been used for Patch-seq mapping (Hodge, Bakken, et al 2019; Ref [7]) and transferred labels from the more recent SEA-AD study [21], which includes approximately 100-fold more cells. Reassuringly, we find good correspondence between original cluster assignments and mapped cell types, with the great majority of cells from the same cluster mapping to one or very few SEA-AD types (**Figure 2B**). Areas of the most disagreement match expectations, with (i) additional resolution in several undersampled inhibitory (e.g., SST) and deep-layer excitatory (e.g., L5/6 NP) types, (ii) particular confusion in gradient cell types (e.g., L2/3 IT cells, Pvalb interneurons), and (iii) lack of representation for most types specifically found in age and AD (cell types with-SEA-AD suffix). As with the other studies discussed in use cases 1 and 2, these studies used different experimental methods, and together validate the utility of mapping of this workflow of applying MapMyCells followed by ACE for cell type annotation and visualization.

We next applied this workflow to two studies of disease whose data was generated from non-author institutions to better represent data from independent users. The first study includes five data sets from the Aligning Science Across Parkinson’s (ASAP) initiative focused largely on donors with different types of PD and including healthy controls. Data from these studies were preprocessed and integrated by ASAP, and then shared with the Allen Institute where cells were mapped to the SEA-AD MTG and whole human brain taxonomies using MapMyCells, and finally provided publicly on the ABC Atlas for community use. Part of this process involved clustering the integrated space; however, no informative cell type names are currently provided for these clusters. To aid in cell type annotation, we include this information as a prebuilt table in ACE (“Parkinson’s disease (ASAP-PMDBS project)”). By exploring individual annotations, we can quickly and easily align most clusters to known cell types; for example, clusters 4 and 1 correspond to Sst and Pvalb interneurons and L2/3 IT cells in neocortex, respectively (**Figure 2C**; left panels), while clusters 25 and 29 correspond to medium spiny neurons and a subset of amygdala excitatory neurons, respectively (**Figure 2C**; right panels). The second study is a multi-omics resource from the PsychENCODE Consortium called brainSCOPE focused on DFC in humans with multiple disease conditions [36]. Seven data sets spanning four brain diseases are publicly available and included as a prebuilt table in ACE, again including initial study data and mapping results to the SEA-AD MTG cell type taxonomy. Overall we find strong alignment between brainSCOPE subclass assignments and SEA-AD subclass assignments, particularly for cells with high mapping confidence (**Figure 2D**;Cramer’s V of 0.861 vs. 0.374 in cells with mapping confidence >0.99 vs. <0.99, respectively). We also identify a type of oligodendrocyte (Oligo_3) which shows significantly lower average donor age than other oligodendrocyte cell types (Kruskal-Wallis, p = 1.5e-111), an effect which does not obviously appear to be associated with disease status (**Figure 2E**), although this could be a good hypothesis for future study.

### Use Case 2: Comparing brain cell types in AD across 11 studies

Multiple published snRNA-seq studies describe cellular and molecular changes in AD [22–31], but they all use different names for neuronal and glial types, making cross-study comparisons challenging. In a previous effort [21], we harmonized snRNA-seq data and associated donor metadata across 11 such studies of AD including SEA-AD and 10 community-based data sets. This involved reprocessing raw sequencing data though the SEA-AD analysis pipeline and then mapping cells to SEA-AD cell types. Here we extend this work by linking mapping results with cell type assignments and changes in abundance with AD reported in the original studies (see above and **Supplemental Figure 2**), saving the results as a pre-built table in ACE called “Alzheimer’s disease (SEA-AD vs. community)”. Assigned cell types for these AD studies match the mapped cell types from the two disease studies discussed above. This standardized cell type naming allows for direct and meaningful comparisons across different neurodegenerative and neuropsychiatric diseases, utilizing the methods and resources we’ve provided.

One key finding from our previous work is that SEA-AD and the two largest studies of DFC [22, 31] show common abundance changes in AD, including loss of supragranular somatostatin (SST) neurons and gain of disease-associated microglia [21]. We used ACE to explore abundance changes with AD across the remaining studies of AD, which were underpowered for direct statistical assessment using an integration approach. These same SST neurons showed consistent cell loss in donors with AD pathology in several other published studies; for example, SST_25 abundances were reported as decreasing with AD six of the nine studies that included neurons (**Figure 3A**, top row). Interestingly, other SST neurons that we identified as relatively spared also were not reported to show decreases with AD in other published studies (e.g., SST_1, **Figure 3A**, bottom row). The only two studies showing decreases in SST_1 abundance in AD are studies that group nearly all SST cells into a single type, suggesting study differences in this case are due to lack of cell type resolution (**Figure 3B**). Changes in abundance of other cell types show less reproducibility between studies. Disease associated microglia (Micro-PVM_3-SEAAD, **Figure 3C**, top row) were reported to show increases in abundance in two other studies, but are not reported as increased in several others, potentially due to lower statistical power [21]. Changes in other glial types are less consistent, with oligodendrocyte and astrocyte types showing decreased and increased abundance with AD in SEA-AD corresponding to types showing a variety of patterns elsewhere (**Figure 3C**, bottom rows).

**Figure 3:**
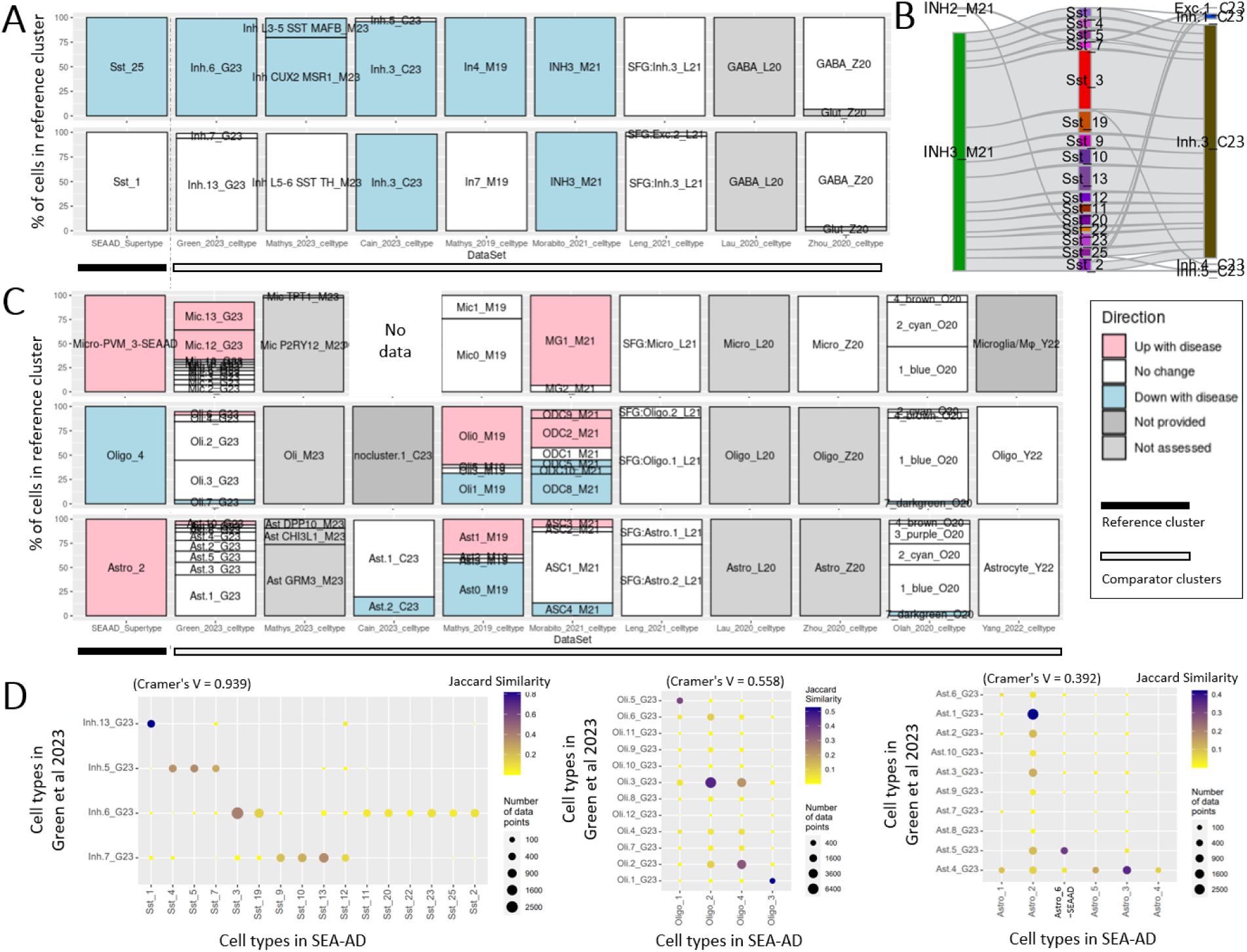
Comparison of cell types across multiple studies of AD. **A)** Cell type assignments in eight other studies of AD for neuronal cells assigned to Sst_25 (top) and Sst_1 (bottom) in SEAAD. Green_2023, Mathys_2023, Mathys_2019, and SEAAD show a consistent decrease in abundance in AD for cells in Sst_25, but not Sst_1 (legend in **C**). **B**) River plots comparing Sst cells in SEAAD (middle column), with Morabito_2021 (left) and Cain_2023 (right) shows that both studies have substantially lower resolution, with most Sst cells mapping to a single type. **C**) Cell type assignments in ten other studies of AD for glial cells assigned to Micro-PVM_3-SEA-AD (top), Oligo_4 (middle), and Astro_2 (bottom) in SEAAD. Abundance changes for microglia are consistent across studies, but less robust than changes in Sst_25, while abundance changes in other glial types vary across studies. **D**) Confusion matrices showing cell type alignment between SEA-AD and Green et al 2023 for Sst neurons (left), oligodendrocytes (middle), and astrocytes (right). SEA-AD identifies more neuronal types and fewer glial types than Green et al 2023 and there tends to be less confusion in neuronal cell type assignments (indicated by a higher Cramer’s V statistic in ‘Sst’ types vs. ‘Oligo’ and ‘Astro’ types).

These inconsistencies in glial results may be due to differences in how cell types are defined; SEA-AD defined more neuronal types, whereas other studies (e.g. [31]) defined more glial types, and cell type definitions tend to be better aligned for neuronal than non-neuronal types across studies (**Figure 3D**). These differences could also reflect discrete and more continuous identities among neuronal and non-neuronal cells, respectively. With more continuous identities in non-neuronal cells, there are a greater number of plausible boundaries that are reasonable, which is influenced by decisions in dataset pre-processing, on which genes are considered through feature selection, on whether and how donor and library effects are modeled, by what clustering parameters are used, and on whether orthogonal experimental data were considered. Together these results highlight the importance both of aligning on a consistent cell type nomenclature when defining cell types and of having tools for visualizing cross-study patterns when studying abundance changes in AD.

### Use Case 3: Translating between reference brain cell type taxonomies

The Allen Institute has developed multiple high-quality and well-annotated cell type taxonomies [4–11, 19, 21] over the past decade, starting with focused studies in individual brain areas and gradually increasing in scope as technologies have improved. Most of these studies are accompanied by web applications on Allen Brain Map that are tied to specific cell type taxonomies. A major challenge is linking cell type names and associated knowledge from earlier studies to more recent studies spanning the whole brain both for scientists within the Allen Institute and for the neuroscience community at large. In particular, we receive many questions asking how to link cell types from our initial, widely-used neocortical studies [7, 8] to more recent studies of the same brain regions that include more data (human) or span the whole brain (mouse and human). A complete list of transcriptomics brain atlases created by the Allen Institute with associated data exploration tools is shown in **Table 1**.

**Table 1:** List of transcriptomics atlases on Allen Brain Map and their associated tools. An extended version available on Allen Brain Map at https://portal.brain-map.org/help-and-community/guide-cell-types. Abbreviations: VISp, primary visual cortex; ALM, anterior lateral motor cortex; MTG, middle temporal gyrus; M1 and MOp, primary motor cortex. Notes: **(1)**, The data exploration tool presents a draft version of the “cross-areal” taxonomy, which does not match the one in the manuscript or in ACE; **(2)** This study has two instances of Transcriptomics Explorer; only one is shown; **(3)** “M1 - 10x genomics (2020)” is a subset of the data from this study; **(4)** Only a subset of SEA-AD tools are presented here. See sea-ad.org for additional information; **(5)** Two separate studies are included in the ABC Atlas that use the same taxonomy generated from Yao et al 2023; **(6)** Data from this study also available on GitHub and at CELLxGENE. EEL FISH is also used in this study but is not available on Allen Brain Map. All tools in this table can be accessed from brain-map.org under the “Cell Types” tab. **(7)** The full name is “ASAP Human Postmortem-Derived Brain Sequencing Collection.”

| Name on Allen Brain Map | Reference | Omics technique(s) | Brain region(s) targeted | Species | Data exploration tool(s) | Note |
| --- | --- | --- | --- | --- | --- | --- |
| V1 & ALM - SMART-seq (2018) | [8] | SMART-Seq v4 | VISp; ALM | Mouse | RNA-Seq Data Navigator: Mouse |  |
| MTG - SMART-seq (2018) | [7] | SMART-Seq v4 | MTG | Human | RNA-Seq Data Navigator: Human |  |
| Multiple Cortical Areas - SMART-seq (2019) | [9] | SMART-Seq v4; 10xv2 | Eight neocortical brain regions (including MTG and M1) | Human | Transcriptomics Explorer | 1 |
| Whole Cortex & Hippocampus (2019/2020) | [5] | SMART-Seq v4; 10xv3 | Isocortex and hippocampal formation | Mouse | Transcriptomics Explorer; Taxonomy browser tool | 2 |
| M1 - 10x genomics (2020) | [6] | 10xv3 | M1 | Human | Transcriptomics Explorer |  |
| Cell Type Knowledge Explorer | [6, 19] | SMART-Seq v4; 10xv3 | M1; MOp | Human; Mouse; Marmoset | Cell Type Knowledge Explorer | 3 |
| Seattle Alzheimer's Disease Brain Cell Atlas (SEA-AD) | [21] | 10xv3; MERFISH | MTG | Human | Transcriptomics Explorer; Allen Brain Cell Atlas; MapMyCells | 4 |
| Mouse whole-brain transcriptomic cell type atlas | [4, 39] | 10xv2; 10xv3; MERFISH | Whole brain | Mouse | Allen Brain Cell Atlas; MapMyCells | 5 |
| Transcriptomic diversity of cell types in adult human brain | [11] | 10xv3 | Whole brain | Human | Allen Brain Cell Atlas; MapMyCells | 6 |
| Cellular and molecular characterization of the aged mouse brain | [20] | 10xv3 | Whole brain | Mouse | Allen Brain Cell Atlas |  |
| ASAP-PMDBS (Parkinson's disease) | (none) | 10xv3 | Multiple brain areas | Human | Allen Brain Cell Atlas | 7 |
| Consensus basal ganglia taxonomy | (none) | 10xv3 | Basal Ganglia (BG) | Human; Macaque; Marmoset | MapMyCells |  |

To facilitate translation of cell type names and (to some extent) knowledge between studies, ACE includes several annotation tables translating cell types between brain atlases, which can be accessed by choosing “Mouse Cell Type Classification” or “Human Cell Type Classification” as the data type in ACE. This section tackles several questions involving translation between reference cell type taxonomies that can be addressed using ACE.

### Which cell types in the whole mouse brain correspond to a specific cell type in mouse whole cortex & hippocampus, and what do we know about them?

By selecting the appropriate data set and exploring individual annotations for the cell type of interest, we see that “059_L6 IT CTX Osr1” includes cells from multiple cell types in the whole mouse brain (**Figure 4A**), suggesting that L6 IT cells are subdivided differently in both taxonomies and that there is higher cell type resolution in the whole mouse brain taxonomy.

**Figure 4:**
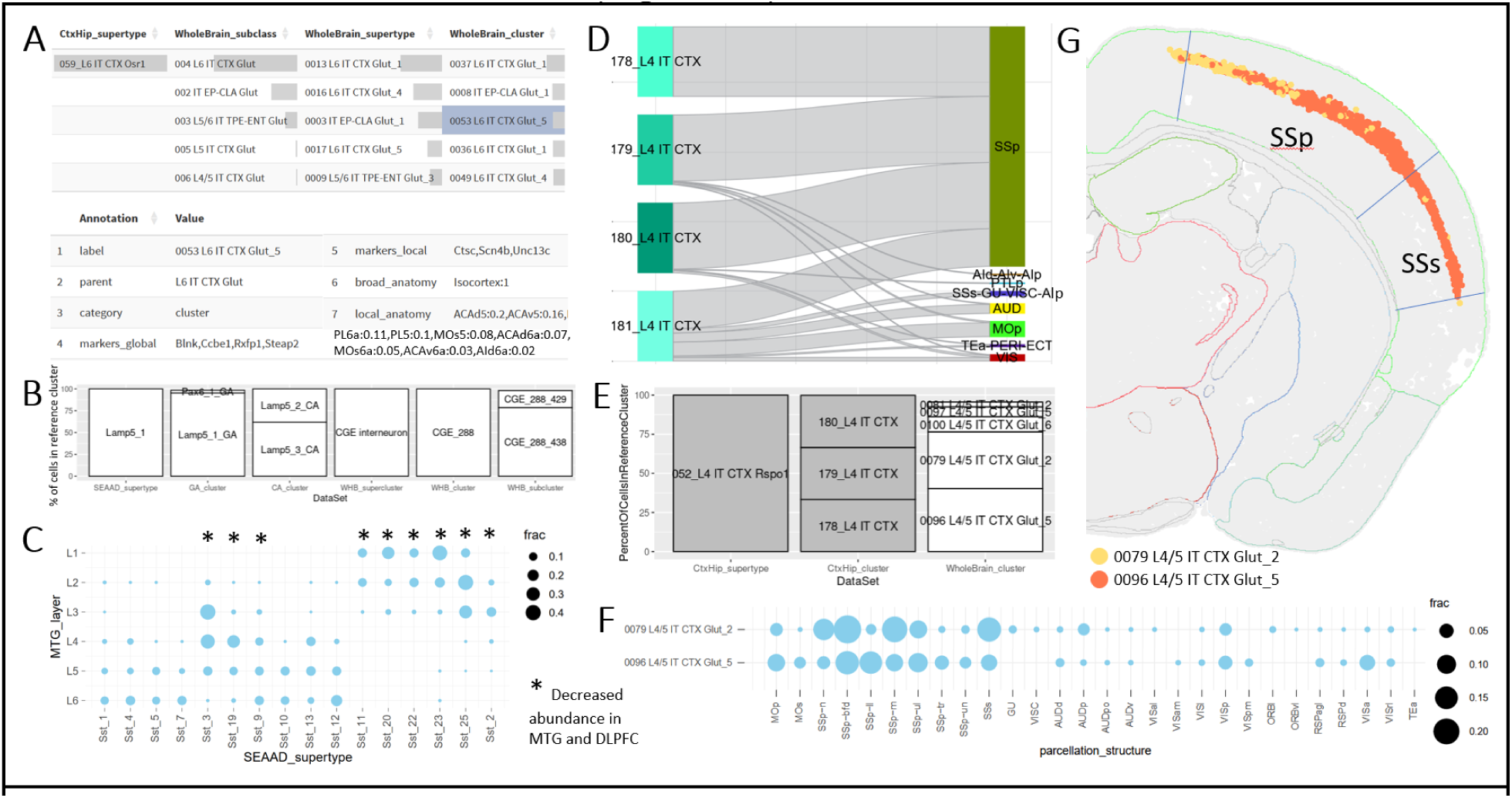
Addressing biological questions through annotation comparison in the healthy adult brain. **A)** Translation of mouse cortical cell types between studies. Top: Bar plot shows annotations of 059_L6 IT CTX Osr1 cells from a study of mouse isocortex and hippocampus (Yao et al 2021) in a newer study of whole mouse brain (Yao et al 2023). Bottom: Cell type annotations for 0053 L6 IT CTX Glut_5 include information about cell type markers and anatomical localization. **B**) Translation of an interneuron cell type (Lamp5_1) across several studies of adult human neocortex or whole brain. **C**) Confusion matrix comparing dissected cortical layer and mapped SEA-AD supertypes (from Figure 3B) for Sst cell from Hodge, Bakken, et al 2019. Dots represent fractions of cells from a cortical layer mapping to a specific SEAAD_supertype (rows sum to 1). Supertypes showing decreased abundance in MTG and DFC in Gabitto, Travaglini et al 2024 (*) were preferentially located in superficial layers. **D-G**) Identification of neurons somatosensory (SS)-specific neuronal cell types. **D**) River plot showing the cluster assignment (left) and anatomic dissection (right) for cells in the only supertype primarily including cells from SSp (052_L4 IT CTX Rspo1). **E**) Bar plot showing the relationship between clusters in whole mouse brain and the three SSp-specific clusters in mouse isocortex plus hippocampus identified in **D**. **F**) Spatial localization in whole mouse brain (x-axis) for the two dominant whole mouse brain clusters identified in **E** (y-axis). Dots scaled as in **C** (rows sum to 1). **G**) A single hemisphere of MERFISH tissue section C57BL6J-63885.43 from the Allen Brain Cell Atlas shows that all cells from these two clusters are highly localized to layer 4 in primary (SSp) and secondary (SSs) somatosensory cortex. Note that panel **G** was generated with the Allen Brain Cell Atlas and not ACE.

Most of the names include the term “L6 IT” suggesting that a consistent broader cell type resolution is retained. Finally, by selecting a particular whole brain cluster, we can identify relevant marker genes and we see that these cells come from multiple neocortical areas, suggesting this cell type is broadly found across neocortex and not restricted to a single brain region. Similar questions can be asked when linking other data sets, including human studies focused on MTG and the whole brain (**Figure 4B**).

### How do we know that somatostatin interneurons impacted in AD are primarily found in superficial cortical layers?

As mentioned above, only a subset of SST types show decreased abundance in AD (**Figure 3A**). We reported previously that these cell types are primarily found in layers 2 and 3 of the human neocortex [21], but how can we use ACE to confirm? Our initial MTG study contained laminar dissections, which are included in ACE. In Use Case #1, we mapped cells from this study to SEA-AD using MapMyCells (**Figure 2B**) and have shared this as a precomputed table in ACE. We find that nearly all SST cells collected from cortical layer 1, 2, and 3 and some SST cells from layer 4 mapped to cell types showing decreased abundance in AD (**Figure 4C**), as reported.

### Are area-specific isocortical cell types also found in subcortical structures and/or spatially organized?

Primary sensory cortices are highly specialized in mammals, with humans having a greatly expanded visual cortex and mice having specialized barrel fields in primary somatosensory cortex (SSp). In humans, several specialized neuronal types are found in VISp that are primarily localized to layer 4 [9]. Using ACE we assessed whether mice showed similarly specialized cell types in SSp. To do this we first plotted a confusion matrix comparing the cell types in mouse cortex and hippocampus with the dissections from which they were collected, identifying a single supertype, 052_L4 IT CTX Rspo1, predominantly found in SSp (**Supplemental Figure 3**). This supertype includes four clusters, three of which are almost exclusive to SSp, and one with mixed expression across other largely primary sensory areas (**Figure 4D**). We then filtered our analysis to only include cells in these three SSp-specific types, finding >75% of these cells were reassigned to two clusters in the whole mouse brain (**Figure 4E**).

We next shifted annotation tables in ACE to one focused on spatial localization of brain cell types and filtered cells to include only those found in 0079 L4/5 IT CTX Glut_2 or 0096 L4/5 IT CTX Glut_5. Consistent with matched clusters from mouse cortex plus hippocampus the majority of cells in these clusters were found in primary (SSp) or secondary (SSs) somatosensory cortex, although the distribution of these two clusters across subcompartments appeared slightly different (**Figure 4F**). To identify the precise spatial localization of cells from these two clusters in the whole mouse brain, we exited ACE and viewed a mouse MERFISH data set with reconstructed spatial coordinates using the Allen Brain Cell Atlas. These two clusters were highly spatially localized in layer 4 of isocortex, extending just beyond the reported boundaries of SSp and SSs, with cells in 0079 L4/5 IT CTX Glut_2 tending to be found more dorsally than cells in 0096 L4/5 IT CTX Glut_5 (**Figure 4G**). Out of the ∼35,000 cells in these two clusters, ∼70% were in SSp, ∼15% in SSs, and only five were found outside isocortex (likely representing errors in label transfer). To assess potential functional relevance of these cell types in somatosensory cortex we used ACE to explore reported marker genes for these clusters [4]. Interestingly, a conditional knockout in rhombomeres 3-5 of the top local marker gene for 0079 L4/5 IT CTX Glut_2 (*Robo3*) has been reported to induce bilateral innervation of the somatosensory thalamus and cortex, resulting in a barrel field of smaller, duplicated barrels representing both functional contra- and ipsilateral sensory inputs, with a decrease in morphological complexity of layer 4 spiny stellate cells within barrels [46]. Thus, the distinct localization of these SSp-selective cell types likely has structural and functional consequences when key genes are disrupted.

These use cases represent common examples of how ACE can be used in cell type classification and comparison across studies; however, the visualizations included are highly generic. As a proof of principle of the versatility of ACE we present two additional use cases focused on disease diagnostics and treatment (**Supplemental Figure 4**) and on exploration of US Census data (**Supplemental Figure 5**) for interested readers (**Supplemental Text 2**).

## Discussion

ACE is an interactive tool that helps researchers compare cell type assignments and other metadata across different datasets. It allows users to upload their own data or choose from several predefined datasets, filter the data to focus on specific data points of interest, and then explore the relationships between the annotations using a variety of interactive visualizations. For example, ACE can show how the classification of a particular cell type changes between two different studies, how user-provided clusters relate to MapMyCells mapping results, or how the abundance of a cell type changes across different disease states. To facilitate usability, ACE includes an extensive user guide (**Supplemental Text 1**) and multiple videos talking through specific ACE features and use cases, along with access to direct user support and link outs to related tools (e.g., MapMyCells). Overall, ACE is a versatile tool that can be used to explore and compare a wide variety of data annotations.

ACE is, to our knowledge, unique in its focus on cell annotations and metadata, its combination of user-provided and public data, and its ease of use, and therefore has a different set of advantages and limitation from transcriptomics viewers like CZI CellXGene [47], Allen Brain Cell Atlas, Cytosplore [48], and Cirrocumulus [49] whose main purpose is gene expression exploration. These transcriptomics viewers excel at visualizing and analyzing single-cell (or spatial) transcriptomics data from a single study, displaying cells in 2D (or in some cases 3D) and allowing color-coding by individual metadata fields. Some also offer features like differential expression calculations and latent space regeneration, and can link cell type knowledge across multiple studies and organ systems; however, they often involve very large files (several gigabytes or more) that can be difficult to work with and require specialized knowledge to create. In contrast, ACE excels in the visualization and comparison of multiple metadata fields from one or more studies, requiring only simple text files and a web browser to do so, making it easily accessible to researchers in any field. However, ACE only includes limited functionality for viewing spatial transcriptomics data, and input files typically do not include gene expression values at all. Although many of ACE’s functions could be replicated with R or Python code, doing so requires moderate coding skills, potentially restricting access for non-computational users. In parallel to how CellXGene includes multiple public sc/snRNA-seq for community exploration, ACE includes pre-built annotation files that link Allen Institute cell type taxonomies to one another and to multiple studies of disease. This feature addresses common requests from both the Allen Institute and the broader neuroscience community, thus allowing ACE to fill a crucial gap in the web tools available on Allen Brain Map. At the same time, ACE can be applied to a broad range of use cases, from cell type characterization to exploration of US Census data. Altogether, ACE represents a complementary tool to widely used transcriptomics viewers for exploration of cell type taxonomies.

ACE has some limitations and opportunities for future improvement. First, since it is a lightweight tool written in R shiny and hosted on shinyapps.io, there are some constraints on file size; more explicitly, ACE becomes noticeably slower when the number of rows x number of columns of columns of the input matrix exceeds a few million (the exact number depends on the number of unique annotation values and the specific visualization), requiring subsampling of large data sets. We mitigate potential biases in our statistical analysis by evenly subsampling by cluster to the extent possible to avoid excluding rare cell types. Second, the ACE code base has been developed over several years to target an evolving set of use cases and uses some out of date R libraries and file formats. This is currently addressed by using a fixed R environment with rig, renv, and a hosted rbokeh instance, and by imposing minimal controlled vocabulary in the cell annotation table that is explained in detail in the user guide (**Supplemental Text 1**). Third, the cell typing field is ever evolving, and therefore ACE will never contain all brain cell type classification studies. However, by allowing upload of user data and by periodically adding additional prepopulated annotation tables to match cell type taxonomies included on Allen Brain Map, ACE will remain relevant moving forward. Fourth, while we are highly confident in all cell type assignments published and hosted by the Allen Institute, there is no guarantee that results from mapping algorithms or comparisons between external studies or user-provided data sets will match to the same extent. In the situation where ACE visualizations and statistics demonstrate discordant results, we would encourage users to assess quality of experimental protocols, filtering of low quality data, and accuracy of computational pipelines, including creation of annotation tables. Finally, while we’ve attempted to anticipate the features that would be of use to the general neuroscience community, there are still outstanding issues to address that we know about (e.g., addition of bookmarking or ‘permalinks’), and likely several to which we are unaware. We plan to continue improving ACE functionality and would encourage anyone that uses the tool to provide feedback on what would be of most value.

### Conclusions

Annotation Comparison Explorer combines standard and custom visualizations into a user-friendly, open-source web tool for exploring categorical and numeric relationships. In particular, ACE can translate cell type classifications and knowledge across studies of health and disease and can relate these annotations to underlying cell metadata such as spatial localization, donor information, and quality control metrics. Please include ACE in your cell type classification studies and provide feedback or contribute to this tool!

## Availability and requirements

*Project name*: Annotation Comparison Explorer

*Project home page*: https://sea-ad.shinyapps.io/ACEapp/

*Operating system(s)*: R Shiny (via shinyapps.io)

*Programming language*: R

*Other requirements*: Web browser

*License*: The 2-Clause BSD License

*Any restrictions to use by non-academics*: License needed

## List of abbreviations

Annotation Comparison Explorer (ACE), Seattle Alzheimer’s Disease Brain Cell Atlas (SEA-AD), quality control (QC), Alzheimer’s Disease (AD), intratelencephalic (IT), extratelencephalic (ET), middle temporal gyrus (MTG), dorsolateral prefrontal cortex (DFC), Sequencing Read Archive (SRA), Parkison’s disease (PD), autism spectrum disorder (ASD), major depressive disorder (MDD), post-traumatic stress disorder (PTSD), peripheral blood mononuclear cells (PBMCs), graphical user interface (GUI), Aligning Science Across Parkinson’s (ASAP), somatostatin (SST), basal ganglia (BG), somatosensory cortex (SS), primary somatosensory cortex (SSp), secondary somatosensory cortex (SSs).

## Declarations

### Ethics approval and consent to participate

All data in this study was either obtained from public sources or stripped of all identifying donor information prior to posting on GitHub.

### Consent for publication

Not applicable.

### Availability of data and materials

ACE is available as a hosted website at https://sea-ad.shinyapps.io/ACEapp/ with source code, associated annotation tables, and scripts for converting primary data sources to annotation tables available on GitHub at https://github.com/AllenInstitute/ACE, in the “R”, “data”, and “data_ingest_scripts” folders, respectively. Direct links to these files, along with locations and references for all primary data sources are included in **Supplemental Table 1**. Although ACE is under active development, a snapshot of the project associated with this manuscript release is stored on Zenodo at https://doi.org/10.5281/zenodo.17382357 and on GitHub.

All figure panels except Figure 2A, Figure 4G, and Supplemental Figure 2 were created using the web version of ACE, either by taking screenshots or by downloading images using the download buttons therein. While some minor rearrangements and additions of supporting information were made for clarity, no edits were made to the data shown in any visualization. Figures 2A and Supplemental Figure 2 were created in PowerPoint. Figure panel 4G was taken as a screenshot from the Allen Brain Cell Atlas, and can be reconstructed using the following URL: https://knowledge.brain-map.org/abcatlas#AQEBSzlKTjIzUDI0S1FDR0s5VTc1QQACSFNZWlBaVzE2NjlVODIxQldZUAADBAFGUzAwRFhWMFQ5UjFYOUZKNFFFAAIAAAFRWTVTOEtNTzVITEpVRjBQMDBLAAIAAAExNUJLNDdEQ0lPRjFTTExVVzlQAAIAAAFDQkdDMFUzMFZWOUpQUjYwVEpVAAICMDA3OSBMNC81IElUIENUWCBHbHV0XzIAMDA5NiBMNC81IElUIENUWCBHbHV0XzUAAAQBAAKEnp2fgjSU5gOCKpijglxyWgQyTlFUSUU3VEFNUDhQUUFITzRQAAWBr6ZKgemsDoGggUeAktXoBgAHAAAFAQFTY24zYgAABgEBAkNCR0MwVTMwVlY5SlBSNjBUSlUAA34AAAAEAAAIVkZPRllQRlFHUktVRFFVWjNGRgAJTFZEQkpBVzhCSTVZU1MxUVVCRwAKAAsBVExPS1dDTDk1UlUwM0Q5UEVURwACNzNHVlREWERFR0UyN00yWEpNVAADAQQBAAIjMDAwMDAwAAPIAQAFAQECIzAwMDAwMAADyAEAAAACAQA%3D.

### Authors’ contributions

KJT aligned all the AD data to SEA-AD cell types. KJT and TL compiled cell metadata from external studies of AD with support from VM. REH compiled cell type knowledge from external studies of AD. JAM developed the software with technical support from AO and BT and with conceptualization and user testing by KJT, TL, REH, and VM. SD developed code for MapMyCells and its alignment with ACE. REH and JAM developed supporting resources for ACE, including the webinar. JAM created all the annotation tables and wrote the manuscript. All authors read and approved the final manuscript.

## Acknowledgements

We wish to thank the Allen Institute founder, Paul G. Allen, for his vision, encouragement, and support. We also wish to thank Lucas Graybuck for providing the initial code base for ACE and to thank Lauren Alfiler for creating both the ACE logo and the Allen Brain Map website “Putting it all together” section used as the basis for **Table 1**.

