## Supplemental Figures 1-5, Text 2 for "Annotation Comparison Explorer (ACE): connecting brain cell types across studies of health and Alzheimer’s Disease"

**Contents:**

Supplemental Text 1-2 (one included herein and one uploaded as a separate file)
Supplemental Tables 1-2 (uploaded as separate files)
Supplemental Figures 1-5 (included herein)

### Supplemental Text

**Supplemental Text 1:** Complete ACE User Guide (also available on GitHub).

*This text document was uploaded as a separate Supplemental File.*

**Supplemental Text 2:** Additional ACE use cases.

This section includes two additional use cases for Annotation Comparison Explorer (ACE) outside of the scope of cell type annotation, presented as proof of principle of the versatility of ACE in comparing any data annotations that can be represented as words or numbers. We emphasize that these two use cases represent exploratory efforts at testing the utility of ACE and would need to be contextualized before any conclusions could be drawn.

*Supplemental Use Case 1: Disease diagnosis and treatment*

One potential utility for ACE is relating disease diagnoses, treatments, and outcomes for different demographics. Here we downloaded information from the California Department of Managed Health Care (DMHC) provided as part of a prior kaggle competition [50] and uploaded it to ACE for exploration. While most diagnoses were relatively evenly distributed between males and females, "OB-Gyn/ Pregnancy" and "Prevention/Good Health" diagnoses were heavily skewed towards females (**Supplementary Figure 4A**), the latter at least partially due to preventive screenings for breast cancer (**Supplementary Figure 4B**). Similarly, several diagnoses differed by age with "Pediatrics" and "Mental" skewing young, while "Central Nervous System (CNS) / Neuromuscular" was most commonly found in the 51–64-year-old group (**Supplementary Figure 4C**). Many of the mental subdiagnoses included neurodevelopmental disorders, while the CNS subdiagnoses included a mix of neurodegenerative disorders and other categories (**Supplementary Figure 4C-D**). Many diagnoses involved only a few treatments, with severe depression being treated with electrical interventions (e.g., brain stimulation), brain tumors requiring neurosurgery and cancer treatment, and paralysis requiring rehabilitation services and durable medical equipment, along with other connections that were less clear to a non-specialist.

*Supplemental Use Case 2: Comparing demographic information across US counties*

As another demonstration of the utility of ACE, we downloaded information about poverty, education, and unemployment rates and household income across counties in the United States collected as part of the U.S. Census [51], joining this information into a single table (**Supplementary Table 2**) for upload to ACE. Overall, metropolitan areas have higher median household incomes than non-metropolitan areas, even after accounting for high variability of income across states (**Supplementary Figure 5A**), with some of the wealthiest areas located near DC, Silicon Valley, and Los Alamos National Labs. We also found county-wide correlations between income and education (as measured by percent of adults with at least a bachelor's education), poverty levels, and (to a lesser degree), unemployment rates (**Supplementary Figure 5B-D**), which all also show differences in metropolitan vs. non-metropolitan areas.

*Supplemental References*

50. Patil P. California Independent Medical Review Dataset. https://www.kaggle.com/datasets/prasad22/ca-independent-medical-review.

51. U.S. Department of Agriculture, Economic Research Service. County-Level Data Sets: Download Data. https://www.ers.usda.gov/data-products/county-level-data-sets/county-level-data-sets-download-data.

### Supplemental Tables

**Supplemental Table 1:** Information and data locations for each prebuilt annotation table available in ACE at the time of manuscript submission.

*This table was uploaded as a separate Supplemental File.*

**Supplemental Table 2:** County level information collected from publicly accessible data at the USDA from https://www.ers.usda.gov/data-products/county-level-data-sets/county-level-data-sets-download-data, and compiled into a single file. The "TableOfCountyInfo" sheet includes the actual data uploaded to ACE in .csv format, while the "VariableDescription" describes each column in the first sheet using definitions from the website above.

*This table was uploaded as a separate Supplemental File.*

### Supplemental Figures

**Supplemental Figure 1: Small example to demonstrate how ACE works. A)** Two sets of annotations for the same set of eight cells; one before and one after a ninth cell is added. Brief definitions shown below. **B)** Encoding of the example from **A** into the cell table (top) and optional annotation table (bottom) required for input into ACE. **C)** Representation of a river plot showing the relationships between the two sets of annotations for the eight cells. Note that ACE will produce a river plot like this one if the tables from **B** are uploaded.

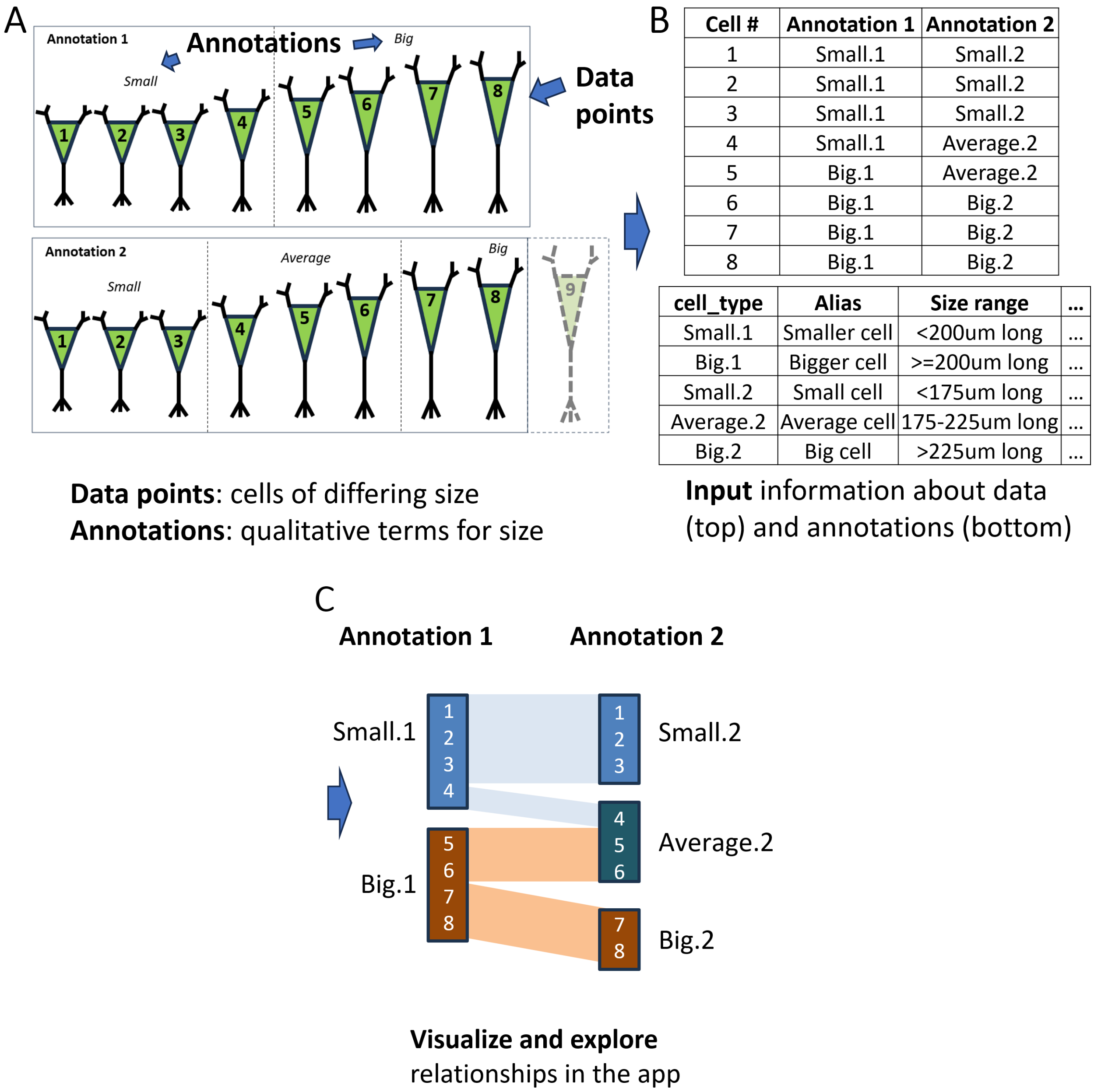

**Supplemental Figure 2: Alignment of 11 studies of AD for inclusion in ACE. A**) Visual representation of showing integration of 11 DFC data sets and transfer of SEA-AD labels to 10 community data sets. **B**) Visual representation showing how SEA-AD cells were assigned community cell types and how changes in AD abundance were recorded.

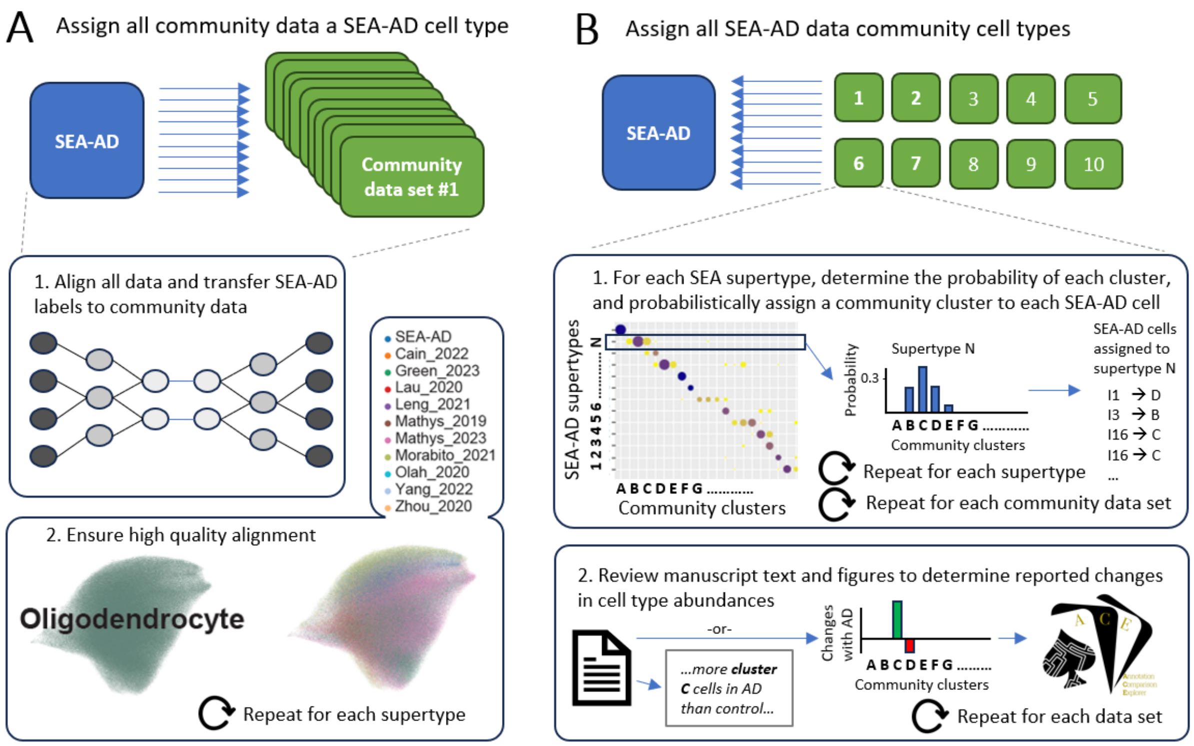

**Supplemental Figure 3: Spatial localization of cell types in mouse cortex and hippocampus.** Confusion matrix showing the dissection structure (x-axis) of cells mapping to each isocortex plus hippocampus supertype (y-axis). Rows are ordered to maximize the diagonal and line up supertypes with similar spatial profiles. Dots scaled as in **Figure 4C** (rows sum to 1). Blue arrow highlights 0053 L6 IT CTX Glut_5, the only cell type primarily collected from SSp. *Note, this supplemental figure spans two pages.*

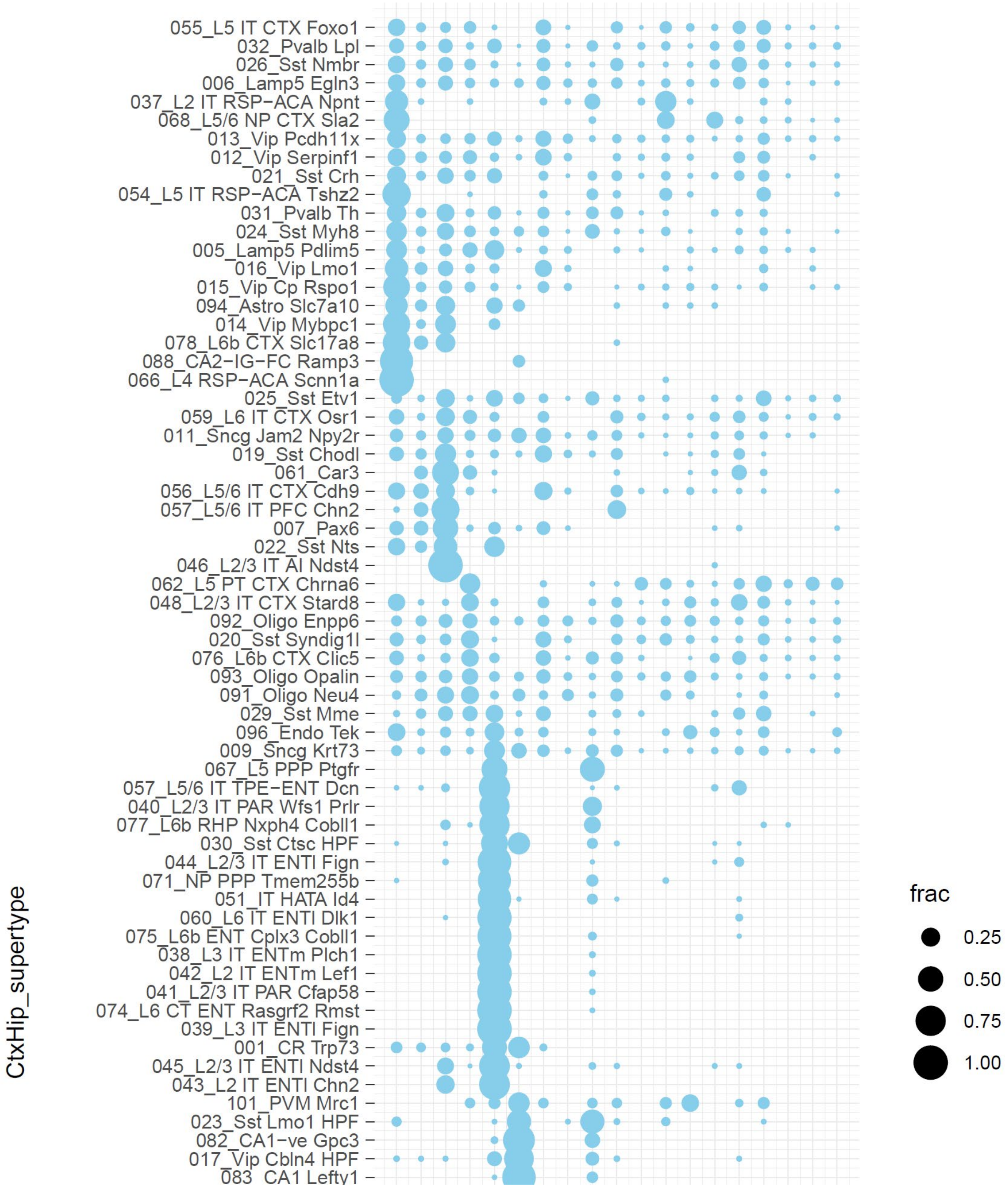

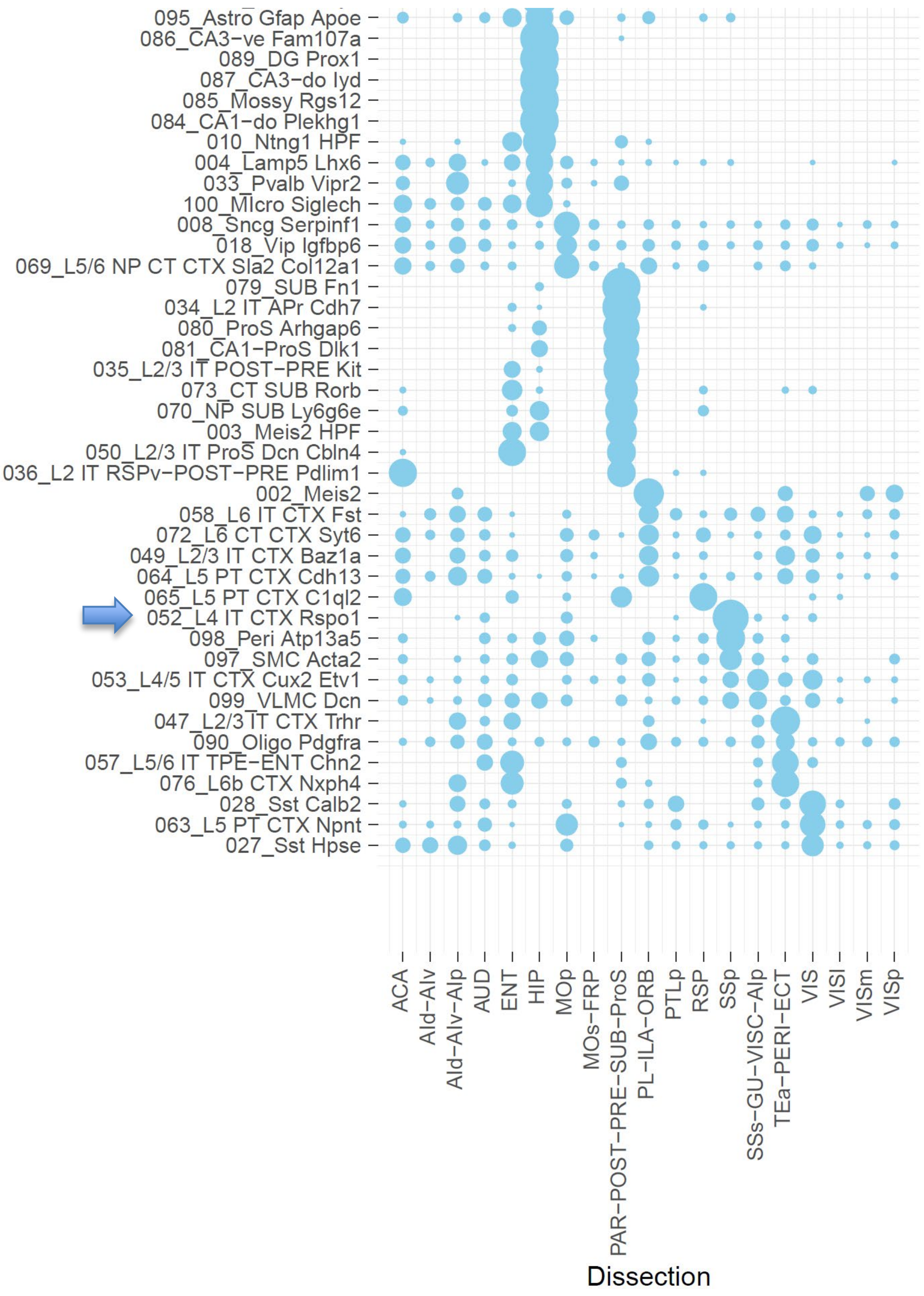
095_Astro Gfap Apoe
086_CA3-ve Fam107a
089_DG Prox1
087_CA3-do Iyd
085_Mossy Rgs12
084_CA1-do Plekhg1
010_Ntng1 HPF
004_Lamp5 Lhx6
033_Pvalb Vipr2
100_Micro Siglech
008_Sncg Serpinf1
018_Vip Igfbp6
069_L5/6 NP CT CTX Sla2 Col12a1
079_SUB Fn1
034_L2 IT APr Cdh7
080_ProS Arhgap6
081_CA1-ProS Dlk1
035_L2/3 IT POST-PRE Kit
073_CT SUB Rorb
070_NP SUB Ly6g6e
003_Meis2 HPF
050_L2/3 IT ProS Dcn Cbln4
036_L2 IT RSPv-POST-PRE Pdlim1
002_Meis2
058_L6 IT CTX Fst
072_L6 CT CTX Syt6
049_L2/3 IT CTX Baz1a
064_L5 PT CTX Cdh13
065_L5 PT CTX C1ql2
052_L4 IT CTX Rspo1
098_Peri Atp13a5
097_SMC Acta2
053_L4/5 IT CTX Cux2 Etv1
099_VLMC Dcn
047_L2/3 IT CTX Trhr
090_Oligo Pdgfra
057_L5/6 IT TPE-ENT Chn2
076_L6b CTX Nxph4
028_Sst Calb2
063_L5 PT CTX Npnt
027_Sst Hpse
ACA
AId-AIv
AId-AIv-AIp
AUD
ENT
HIP
MOp
MOs-FRP
PAR-POST-PRE-SUB-ProS
PL-ILA-ORB
PTLp
RSP
SSp
SSs-GU-VISC-AIp
TEa-PERI-ECT
VIS
VISl
VISm
VISp
Dissection

**Supplementary Figure 4: Disease diagnosis and treatment data. A)** Dot plot showing the fraction of male vs. female individuals with each diagnosis (rows sum to 1). Blue arrow indicates the subset of data points used in panel **B**. **B**) River plot showing the relationship between gender, treatment category, and treatment subcategory for individuals diagnosed with "Prevention/Good Health". **C**) Dot plot showing the fraction of individuals of different ages with each diagnosis. Legend from **A** applies but note the ordering of diagnoses is different. Purple arrows indicate the subset of data points used in panels **D** and **E**. **D-E**) Dot plot showing the relationships between disease subcategory and treatment category for individuals diagnosed with "Mental" (**D**) or "Central Nervous System (CNS) / Neuromuscular" (**E**). Fraction of individuals in each treatment category use the same legend in both panels (rows sum to 1). Similarly, dots are color-coded by treatment category to further allow linking between panels.

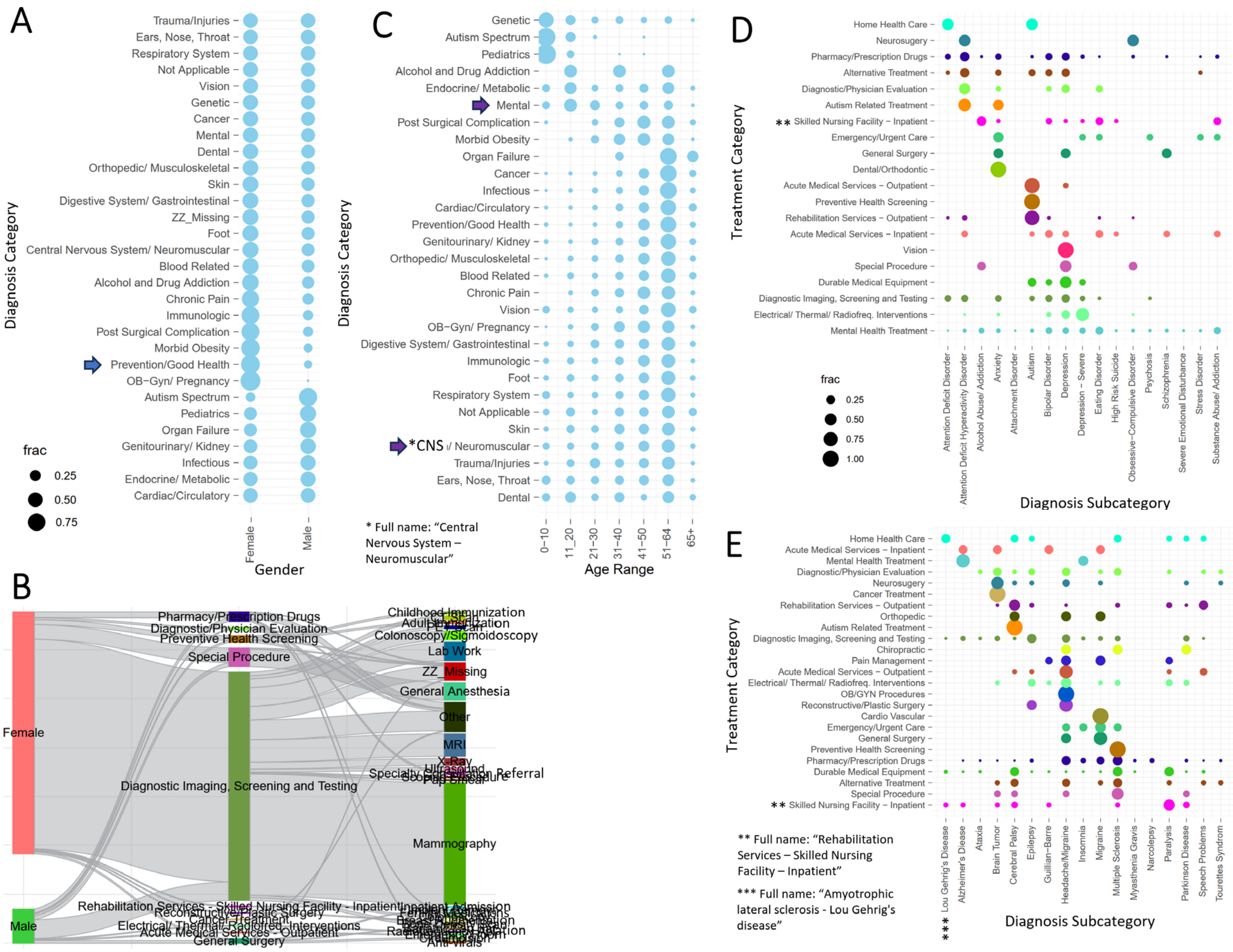

**Supplementary Figure 5: Comparison of US county-level population data. A)** Bee swarm plots showing median household income for all US counties identified as metropolitan (Metro) or non-metropolitan (Non-Metro) area, separating and color-coding by metropolitan designation (left plots) and separated out by state (right plots). A few counties with high median income are highlighted. **B-D**) Scatterplots showing the relationship between correlated metrics and color-coded as in **A**. PCTPOVALL_2021 is the estimated percent of people of all ages in poverty in 2021.

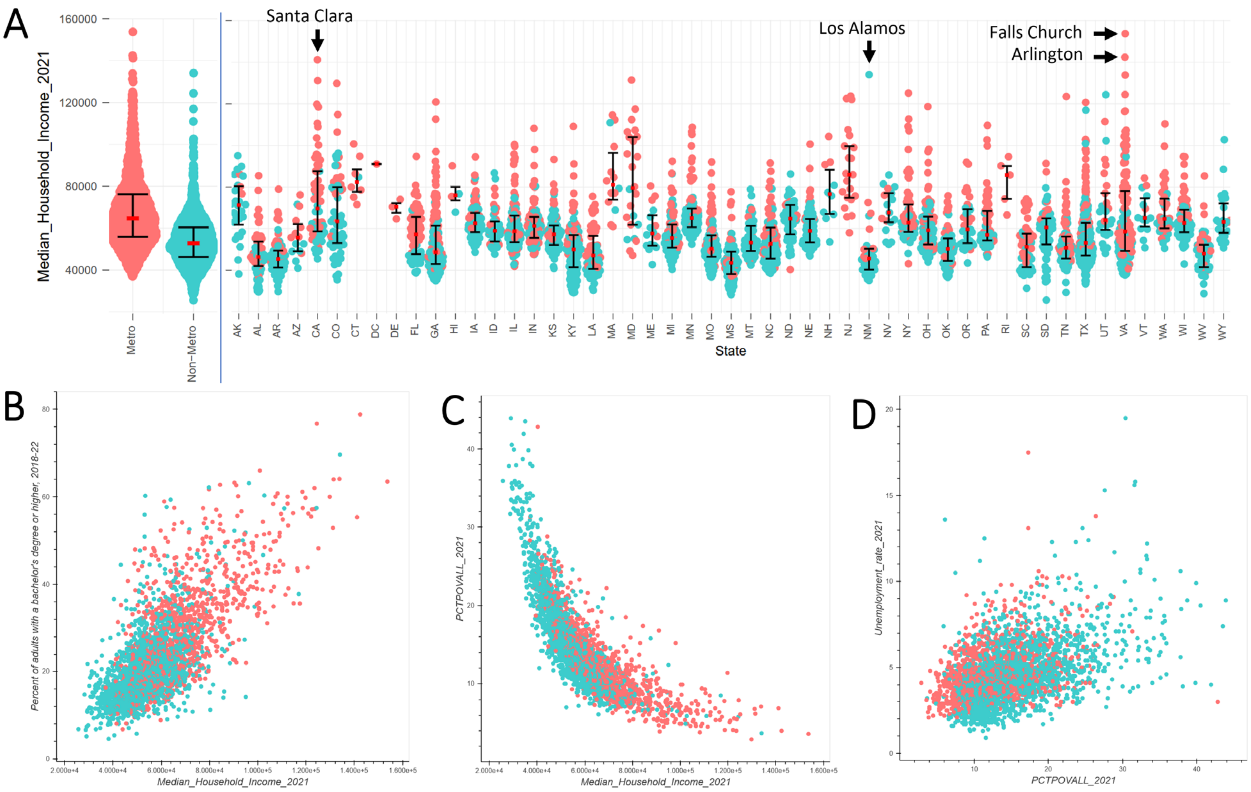
