## Supplemental Text 1 for "Annotation Comparison Explorer (ACE): connecting brain cell types across studies of health and Alzheimer’s Disease"

**ACE is a web (and associated R shiny) application for comparison of different annotations of the same dataset.** ACE was designed (but is not limited to) use cases surrounding comparison of cell type definitions and other cell metadata. Examples include (i) cell type assignments (e.g., from different mapping/clustering algorithms or taxonomies), (ii) donor metadata (e.g., donor, sex, age), and (iii) cell metadata (e.g., anatomic location, QC metrics). Additionally, this tool can compare results across more than two taxonomies, for example for annotating a novel taxonomy with information from multiple existing taxonomies or for comparison of data from multiple studies of Alzheimer's disease.

**Most folks will want to access ACE as a website at https://sea-ad.shinyapps.io/ACEapp/.**

ACE can also be run in RStudio as a shiny app by following the instructions shown at the GitHub repository https://github.com/AllenInstitute/ACE. The codebase is open-source, and we encourage contributions.

Note that ACE is under active development and the interface may not match the content of the User Guide *exactly*. If you run into issues or more generally have any thoughts or opinions about the app, or you would like to contribute to this app, please reach out via the big button on the app or by submitting a GitHub issue.

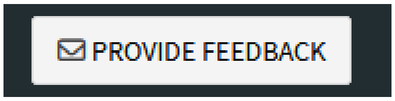


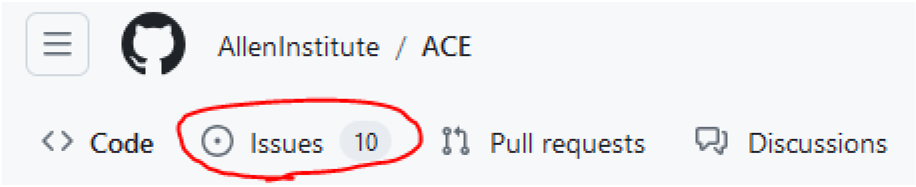


----------

*App developed by Jeremy Miller with support from Aaron Oster and Bosiljka Tasic, using some original code developed by Lucas Graybuck. Included annotation tables were created by Jeremy Miller, Kyle Travaglini, Tain Luquez, Rachel Hostetler, and Vilas Menon. Logo credit: Lauren Alfiler.*

----------------------------

### Use Cases

ACE was created to address a number of use cases not easily addressed using interactive tools.

- **Compare cell type results across published studies of Alzheimer's disease (AD)**: We have assigned cell types from the Seattle Alzheimer's Disease Brain Cell Atlas (SEA-AD) classification to cells from 10 studies of AD and have recorded changes in abundance with AD reported in each study. ACE allows users to interactively explore these relationships and uncover which cell types show common AD-associated changes across studies. (Check out Supertype Sst_25!). Tables aligning cell types from other disease studies (e.g., from PsychENCODE) with Allen Institute taxonomies have also been added (in mid-2025).
- **Convert cell type names between different Allen Institute taxonomies**: Over the past decade the Allen Institute has created multiple cell type classifications in brain, including recent studies in whole mouse and whole human; however, understanding what a specific cell type defined in mouse visual cortex or human middle temporal gyrus in 2018 is now called is not straightforward. ACE includes a variety of translation tables providing easy conversion between these different taxonomies using matched data sets.
- **Compare mapping and clustering results for user-provided data**: MapMyCells provides an easy way for users to assign Allen Institute cell types to user-provided single-cell or spatial transcriptomics data. ACE allows researchers to see how results from their own clustering analysis relate to assigned types for a given set of cells. A separate tutorial for how these two tools can be used in parallel is included on the MapMyCells landing page and is also described in the ACE publication.
- **Identify which cell types are in which brain regions:** Recent publications of cell type classifications in whole mouse and whole human brain have found that many of the several thousand cell types show spatial localization. This extends earlier studies of multiple cortical areas, which found cell types specific to visual cortex and hippocampus, among other brain regions. ACE allows researchers who are focused on a particular brain region to easily filter cells by brain region (or other metadata) and determine which cell types are still present after filtering, or alternatively, to determine the proportion of cells of a given type that are found in different brain structures.
- **Assess data quality by viewing and comparing QC metrics per cluster:** Standard QC metrics, like # of genes detected, % mitochondrial reads, doublet scores, and lab-based pass/fail calls can be extremely useful in determining whether or not clusters defined in a cell type analysis represent "real" biological cell types/states. ACE allows you to plot any numeric values in the context of any categorical variable, so you can quickly scan whether certain clusters have higher or lower QC metrics than most others. The "Explore individual annotations" tab also allows direct comparison of other categorical data to individual clusters (e.g., look how many cell types are included in "cluster_000" from the "Parkinson's disease" study!).
- **Visualize spatial location or morphoelectric features of cells of a given type**: A number of techniques allow a researcher to determine cell type of cells in other contexts. For example, MERFISH provides the specific spatial location of a cell with known type, while Patch-seq allows one to assign a transcriptomic type in a cell with measured morphology and electrophysiology. ACE includes tools for plotting any numeric values in the context of any categorical variable, so you can quickly scan whether certain clusters have higher or lower morphoelectric values. In addition, ACE can compare and color code any numeric values, allowing one to visualize where cells of a given type are located in physical space or in a plot of any two features of interest.

- **Compare the abundance of cells in different situations (e.g., by sex, disease state, hemisphere, or brain region)**: Several glial cell states are known to appear in response to neurological insults, while structural differences in left and right hemisphere may or may not translate to cell type differences. ACE provides a few tools for visualizing such abundance differences through confusion matrices and river plots, or direct inspection of individual metadata fields of cell types.
- **Visualize metadata on a UMAP (or other 2D latent space)**: UMAPs are popular visualizations for exploring gene expression data. While other tools like CZI CellXGene are better suited for this task, if 2D coordinates are provided, ACE can color-code any subset of the cells by any provided metadata, allowing one to see where in the UMAP cells from a given cell type are located, for example.
- **Download plots and data**: Most plots can be downloaded in various sizes for use in your own scientific publications, and the underlying data for many plots is also available if you'd prefer to expand your analysis or plot the data in other ways.
- **Perform statistical analysis**: For many plots a statistical analysis assessing its difference from a random distribution can be performed with a single click. The specific statistic depends on the plot, and data can be downloaded for performing additional statistics.
- **Perform any other metadata comparison**: While ACE was created with cell type analysis in mind, it is ultimately a general tool for visualizing different properties of the same entities, meaning that its uses are only limited by your data and your imagination. Other potential questions ACE could address include:
  - Do more students with red or blond hair also wear glasses?
  - What is the average age of workers in different professions?
  - Physically where in the USA are Republican vs. Democrat-leaning cities located?
  - What is the overlap between the Gyral and modified Brodmann anatomic atlases in human brain?

If you find ACE useful, **please let us know how you use this tool**! Also, if you are having trouble getting ACE working for your use case, please let us know that as well.

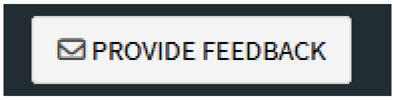

-----------------------------

### How does ACE work?

ACE is an annotation comparison tool. Its input is a single set of data (typically, but not necessarily, cells) with different annotations for those cells (typically, but not necessarily, cell type assignments and/or other cell metadata). The application itself then visualizes these annotation relationships in a variety of ways.

**Example #1: height of adults**

As a simple example, let's assume that we have 11 adults participating in a medical study. They are first asked to measure and self-report their height ("Annotation 1" below), and all but the tallest two individuals comply. Researchers use these data to split these data into short vs. tall. Then a bit later, all 11 adults come into the office and get their heights measured ("Annotation 2"). With the remaining two individuals included, researchers now decide that there should actually be three categories of height: "Short", "Average" and "Tall".

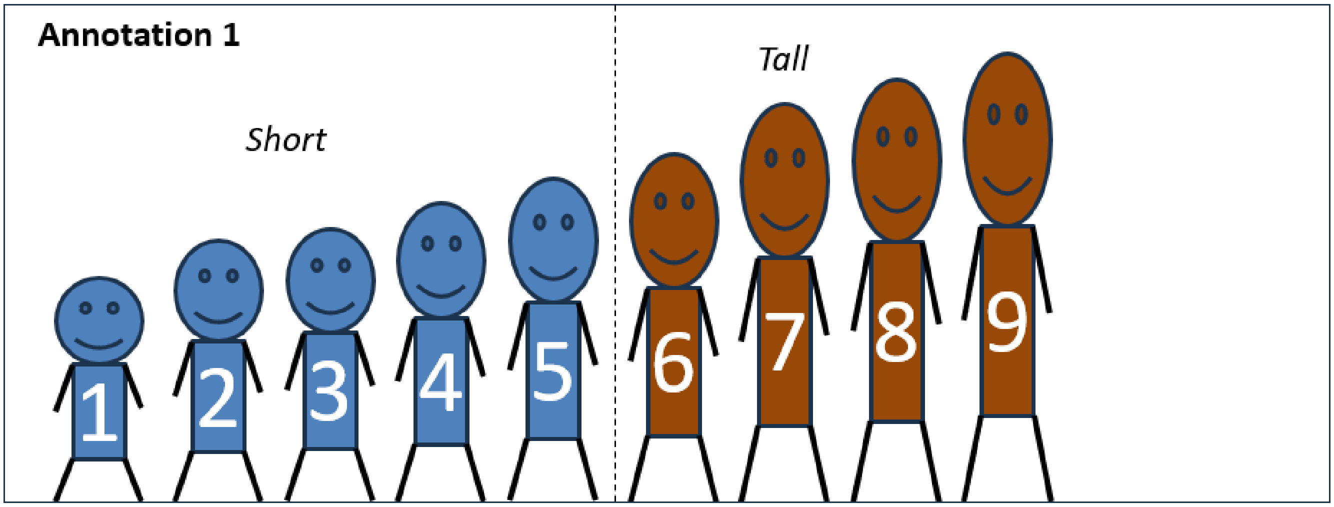


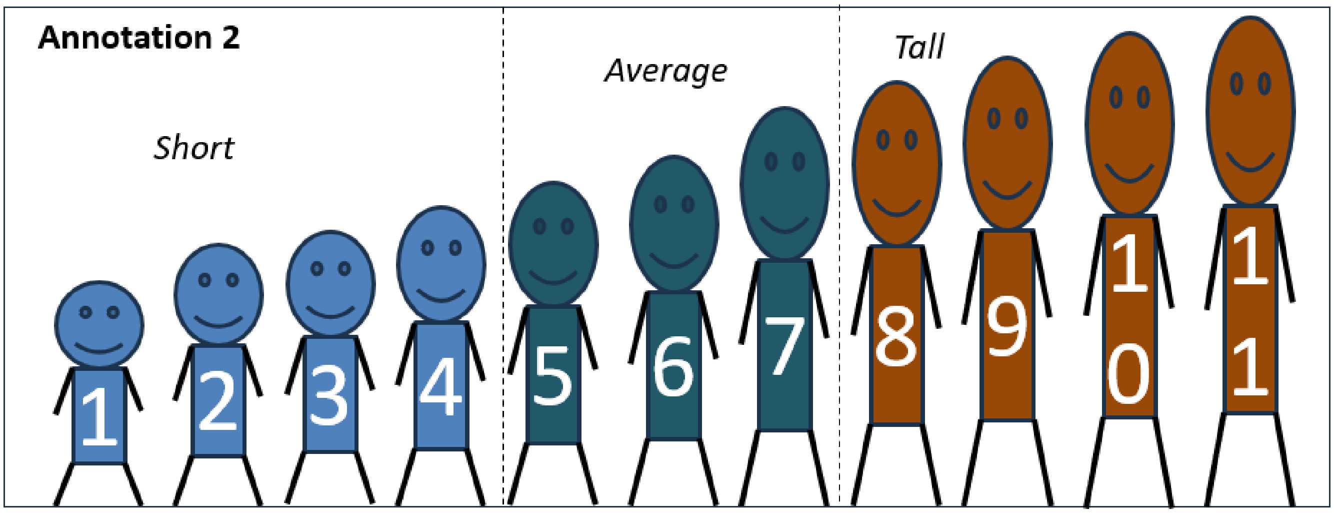


This example mirrors the case when we have a single set of cells that initially had one cell type annotation and later had another cell type annotation, potentially with the addition of more data. Let's pivot to that exact example right now.

**Example #2: Size of cells**

In this analogous example, instead of having people of different height, we now have pyramidal neurons that we'd like to categorize by size. With eight cells an annotator divided these neurons into two groups, but with addition of a ninth cell, another annotated now divided the cells into three groups, causing the initial eight cells to have multiple annotations of their size.

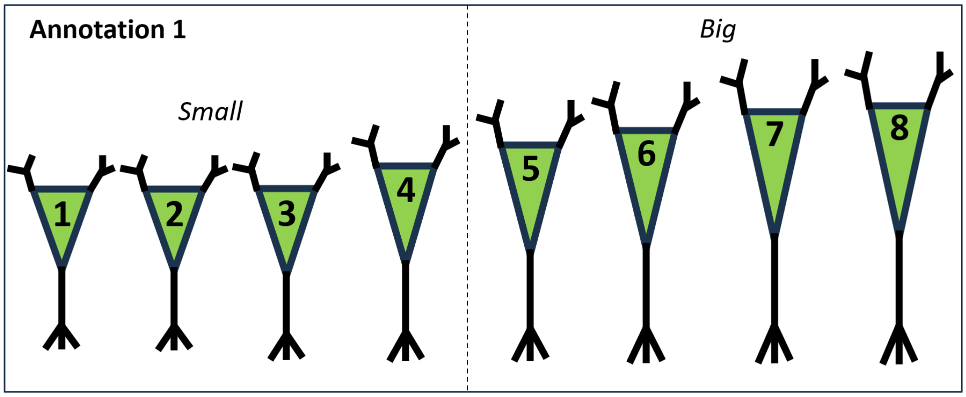


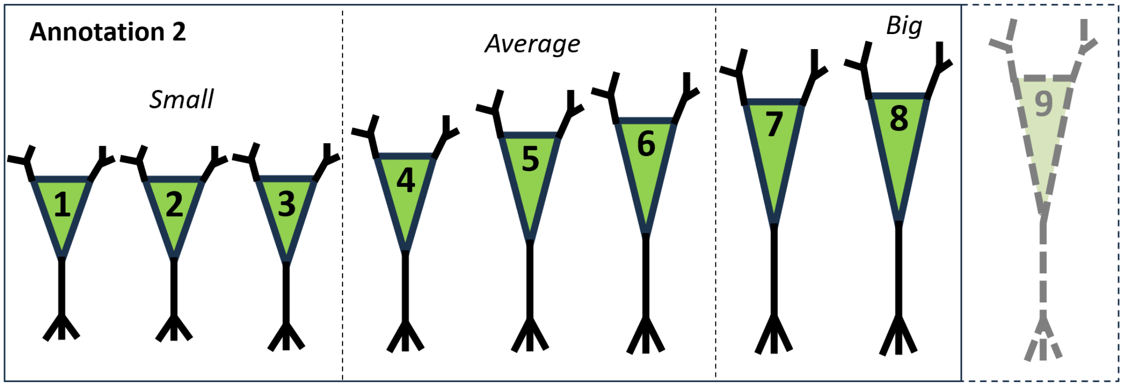


How can we visually and statistically compare these different size annotations? Using ACE of course!

First, we need to prepare the input files. ACE takes as input two files. The first input is the cell annotation file:

| Cell # | Annotation 1 | Annotation 2 |
|---|---|---|
| 1 | Small.1 | Small.2 |
| 2 | Small.1 | Small.2 |
| 3 | Small.1 | Small.2 |
| 4 | Small.1 | Average.2 |
| 5 | Big.1 | Average.2 |
| 6 | Big.1 | Big.2 |
| 7 | Big.1 | Big.2 |
| 8 | Big.1 | Big.2 |

Each row represents a single data point (pyramidal neuron), and each column represents a different annotation (in this case label of cell size). This is essentially a table version of the graphic from above. Because ACE relies on using the same data set, only the eight neurons with both sets of annotations would be used. Neuron #9 (and more generally, any data point with missing annotations) is either ignored by ACE or presented with an annotation indicating missing data.

The second (optional) input file provides additional information about each annotation. In this case there are five values used to describe each neuron's size across the two annotation columns:

| cell_type | Alias | Size range | Annotation overview | Date |
|---|---|---|---|---|
| Small.1 | Smaller cell | <200um long | Jeremy's assessment | 11/23/2024 |
| Big.1 | Bigger cell | >=200um long | Jeremy's assessment | 11/23/2024 |
| Small.2 | Small cell | <175um long | Andrew's assessment | 12/12/2024 |
| Average.2 | Average cell | 175-225um long | Andrew's assessment | 12/12/2024 |
| Big.2 | Big cell | >225um long | Andrew's assessment | 12/12/2024 |

These values can be whatever you want (although currently the column with the metadata values needs to be called "cell_type") and are only used if provided. This allows you to easily review other knowledge of a single metadata value in context, and also provides a default order for annotations (e.g., small < average < big), both of which are helpful when there are hundreds to thousands of values in total.

Once these data are input, ACE provides as number of ways to visualize the comparisons between two (or more) annotations (much more on this below). For example, ACE can make river plots (Sankey diagrams) showing how the cell size changes between two annotation fields. For our example above, it would look like this (the left plot diagrams the details, while the right plot is an actual screenshot from ACE using the above input files):

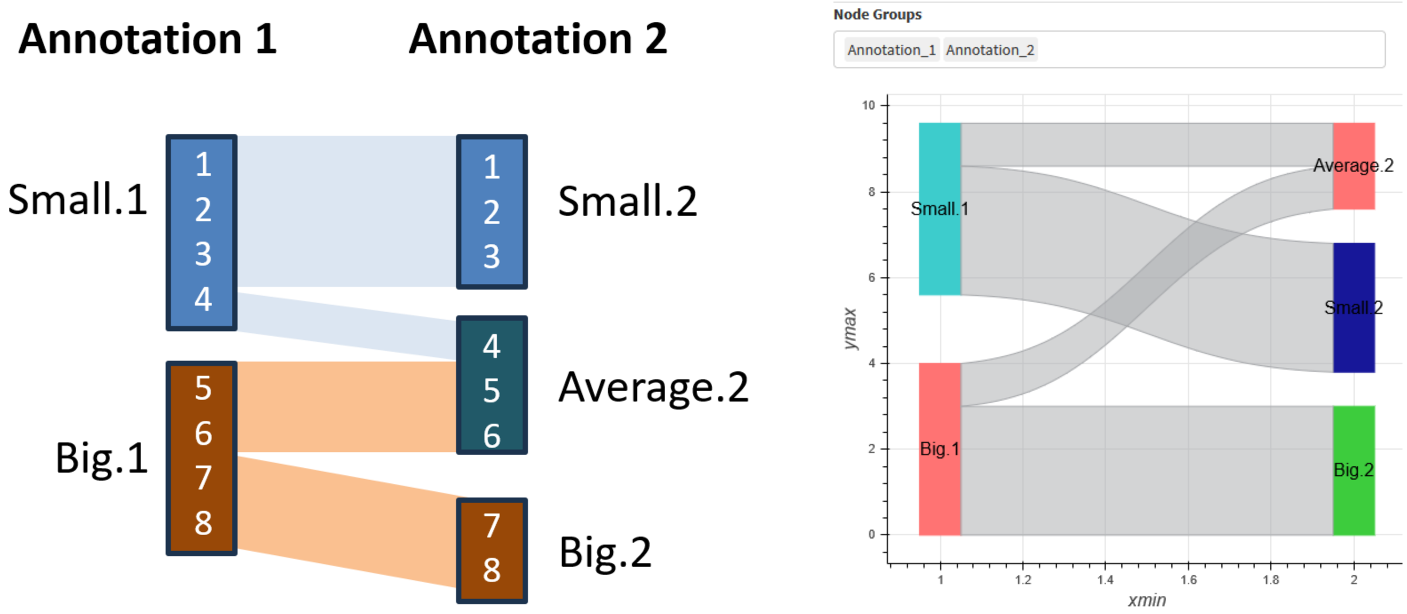


In this case, cells 1-3 are always considered small, 7-8 are always big, and 4-6 change their annotation in a way that can be visualized using the "rivers" between the two columns of colored rectangles. If we look at the metadata table above, we can see that the reason these individuals change height categories is not because they change height, but rather because the definition of what we mean by short or tall has changed between the two annotations (e.g., 'small' changed from <200um to <175um).

ACE provides additional visualization options that are described in detail below, but hopefully this overview provides a general sense for what ACE is doing and how it works. The next section provides many additional use cases of ACE, and the remaining sections go into detail about how the tool itself is used.

-----------------------------

### Getting started

To use ACE, simply go to the website or launch the tool in R (to run locally, see the instructions on GitHub).

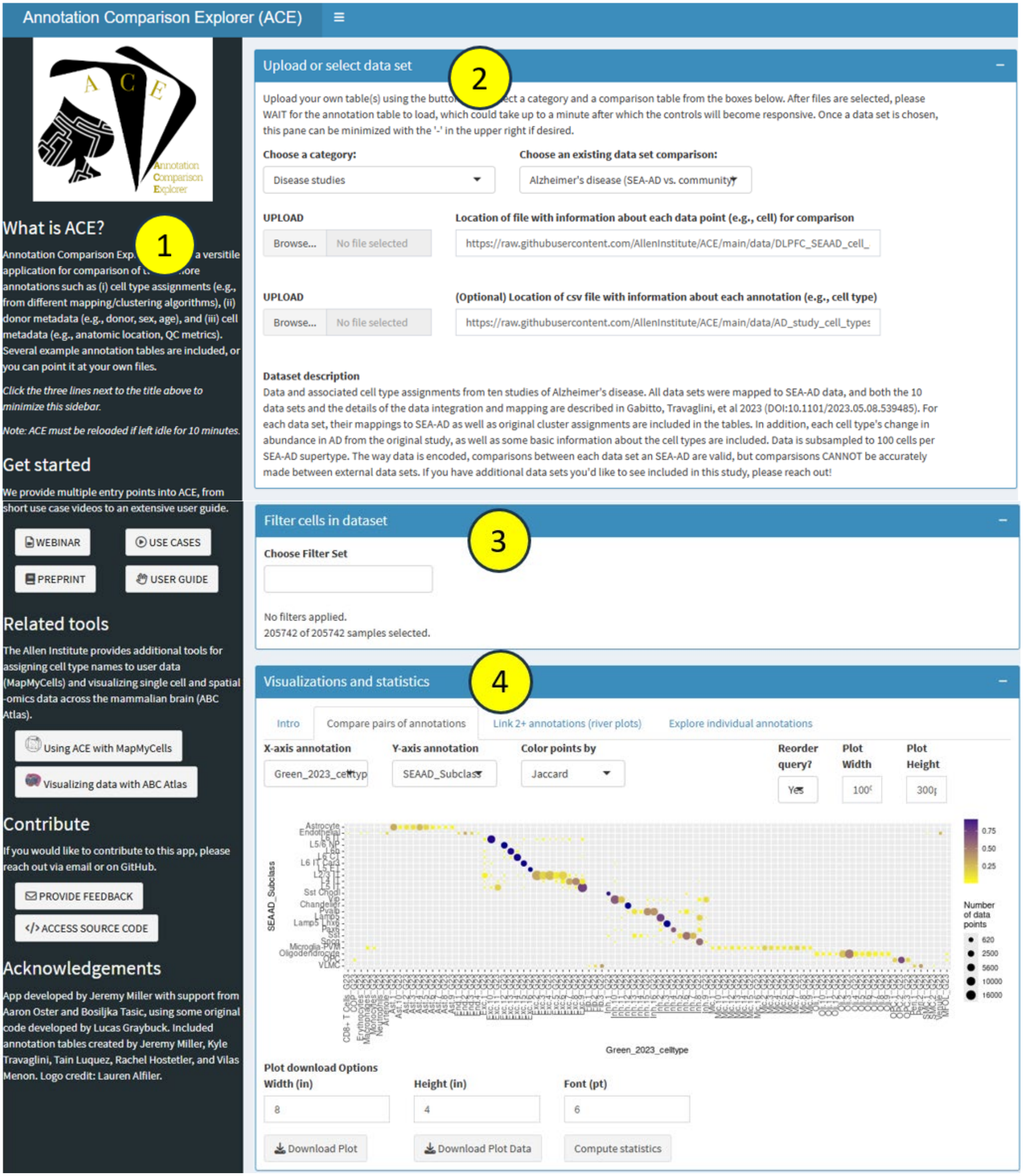

Once open, ACE is an interactive tool that consists of the four panes shown above:

1) **Informational pane**: The first pane contains general information about ACE, including what it is, links out to this guide and other resources to help you use the app, and acknowledgements of app developers. During use, this pane can be minimized by clicking on the three lines at the top of the screen.
2) **Select data set**: This pane allows you to choose a preset annotation table to compare or to upload your own table(s) and is the starting point for all other ACE functionality. As of October 2025, preset annotations fall into four categories:
   a. **Disease studies**: Several tables linking cell type assignments from published community studies of AD, Parkinson's disease (PD), autism spectrum disorder (ASD), post-traumatic stress disorder (PTSD), and more with the SEA-AD cell type taxonomy defined for neocortex.
   b. **Human/NHP cell type classification**: Multiple tables i) translating cell types in middle temporal gyrus (MTG) between cell type taxonomies and with a study of whole brain, ii) exploring cell types in basal ganglia (BG) across species, and iii) describing cell types in human blood.
   c. **Mouse cell type classification**: Multiple tables translating cell types in visual cortex, additional cortical areas, and whole mouse brain between cell type taxonomies, along with another table comparing brain cell types and anatomic structures, and a final table focused on mouse aging.
   d. **Patch-seq (shape + function + genes)**: Tables comparing cell type annotations based on gene expression and multiple modalities for patch-seq cells from five studies and relating morphoelectric features to cell type definitions.

   Please read the "Dataset description" that appears when selecting a comparison table to learn more.
3) **Filter cells in dataset:** After selecting a comparison table, this pane allows you to choose which rows (e.g., cells) to include in the visualizations. *This step is critical when viewing tables with many rows.* For example, one could choose to only look microglial cells, or to focus on cell types in hippocampus. For categorical variables there is an 'invert' feature allowing you to select all cells except a given selection and a box that will add any values matching a specific string. For numeric values a histogram view aids selection of the numeric value range to retain.
4) **Visualizations and statistics**: Once the annotation table and relevant rows are chosen, all the functionality of ACE takes place here. The tabs in this pane provide different tools for comparing cell-level annotations and (optionally) providing additional information about each annotation. Most of this user guide is dedicated to these tools, but a brief overview of them can be found in the "Intro" tab, reproduced here:
   a. **Compare pairs of annotations**: This panel shows different plots depending on the data type. When comparing categorical variables (most cases), a confusion matrix is shown where points can be sized or colored in a variety of ways, including based on Jaccard distance (default). When comparing numeric vs. categorical values a box-and-whiskers + dot plot is shown. An example of this would be in viewing mapping probabilities for each cell type. Comparisons of pairs of numeric annotations are done in a separate panel. For all visualizations in this tab, statistics can be calculated with the push of a button.
   b. **Link 2+ annotations (river plots)**: This panel shows a river plot (also called a 'Sankey plot' or 'Sankey diagram') which chains together relationships between two or more categorical variables. The order variables are listed matters, as the overlap between each pair of adjacent annotations are shown. The second and subsequent node group values are reordered to minimize the number of crisscrossing rivers.
   c. **Explore individual annotations**: This tab focuses on exploring individual annotations (e.g., the "subclass" called "SST"). It includes plots and graphs showing all values of different annotations for a given input annotation, as well as the set of additional metadata for any annotation value selected from the table. Essentially, this is a specialized extended version of a river plot where the data is filtered for a specific metadata value. This is the only place that the "(Optional) Location of csv file with information about each annotation (e.g., cell type)" file is used, aside from the order of cell types, which is also used in the previous panels.
   d. **Compare numeric annotations**: This tab focuses on comparing pairs of numeric annotations. Three common use cases are (1) visualizing 2D UMAP coordinates, (2) showing spatial positions

of cells in a MERFISH tissue section, and (3) comparing morphoelectric features of specific types of cells. Cells are visualized using an interactive scatter plot with hover capabilities. Correlation can be calculated with the push of a button, which is useful for a subset of use cases.

**Note that not all functionalities will be available for every annotation table**. Depending on the types of data provided in the input tool, certain tabs may not work or may be omitted.

----------------------------

### Input data for ACE

At its core, ACE is a method for comparing different cell-level annotations and (optionally) providing additional information about each annotation. Some comparison tables are included, or users can provide their own.

The input to ACE one or two comparison tables:

#### **(1)** Cell data table

The primary input is a **required** table where rows correspond to individual items (typically cells or nuclei) and columns correspond to metadata such as cluster assignments, mapping results, different levels of the taxonomy hierarchy, donor information, or cell QC metrics:

| | A | B | C | D | E | F | G | H |
|---|---|---|---|---|---|---|---|---|
| 1 | **sample_name** | **SEAAD_supertype** | **subclass** | **class** | **GA_cluster** | **CA_cluster** | **sex** | **donor_name** |
| 2 | AAACCCACAACTC | Pax6_1 | Pax6 | GABAergic | Pax6_1 | Pax6_1 | M | H18.30.002 |
| 3 | AAACCCACACGGT | L5/6 NP_1 | L5/6 NP | Glutamatergic | L5/6 NP_1 | L5/6 NP_3 | M | H18.30.002 |
| 4 | AAACCCACACTCT | L5 IT_7 | L5 IT | Glutamatergic | L5 IT_7 | L5 IT_1 | M | H18.30.002 |
| 5 | AAACCCACATCAG | L6 CT_2 | L6 CT | Glutamatergic | L6 CT_2 | L6 CT_1 | M | H18.30.002 |
| 6 | AAACCCAGTGTCG | L4 IT_2 | L4 IT | Glutamatergic | L4 IT_2 | L4 IT_2 | M | H18.30.002 |
| 7 | AAACGAAAGCACT | Astro_1 | Astrocyte | Glia | Astro_1 | Astro_4 | M | H18.30.002 |
| 8 | AAACGAACACAA | L5 IT_2 | L5 IT | Glutamatergic | L5 IT_2 | L5 IT_1 | M | H18.30.002 |
| 9 | AAACGAACACGCT | L2/3 IT_5 | L2/3 IT | Glutamatergic | L2/3 IT_5 | L2/3 IT_3 | M | H18.30.002 |
| 10 | AAACGAAGTACCC | L5 IT_2 | L5 IT | Glutamatergic | L5 IT_2 | L5 IT_1 | M | H18.30.002 |
| 11 | AAACGAAGTTCCG | L5 IT_2 | L5 IT | Glutamatergic | L5 IT_2 | L5 IT_1 | M | H18.30.002 |
| 12 | AAACGAAGTTGCG | Vip_9 | Vip | GABAergic | Vip_8 | Vip_8 | M | H18.30.002 |

User-provided files can either be (1) located on the web (definitely on GitHub, and probably on other URLs but I haven't checked) and provided as a URL in-line or (2) uploaded using the Upload button. Access to local machines is also possible using the R-Shiny version but not on the web version.

Multiple file types are accepted as input:

- A **csv file**, like the one shown above.
- A **gzipped csv file** (e.g., XXXXX.csv.gz)
- An **h5ad file** in the Allen Institute Taxonomy (AIT) format used with scrattch (the obs field is read)—this would likely work for other h5ad files (e.g., input files for CZI CellXGene), but I have not tested it
- A directory containing a **feather** file called 'anno.feather' file with annotations included therein. (This format is likely most useful for Allen Institute employees to visualize taxonomies on molgen-shiny/.)

#### **(2)** Metadata table

The second file is optional, but if provided includes information about each individual piece of metadata. Specifically, each row corresponds to an entry in one of the columns from the cell table (e.g., a specific cluster or subclass) and columns correspond to whatever information about the metadata that you want to share. Here is one example that provides some information about cell sets from SEA-AD:

| 1 | cell_type | level | study | direction | description | notes |
|---|---|---|---|---|---|---|
| 2 | Neuronal: GABAergic | class | SEA-AD | not_assessed | All inhibitory neurons | |
| 3 | Neuronal: Glutamatergic | class | SEA-AD | not_assessed | All excitatory neurons | |
| 4 | Non-neuronal and Non-neural | class | SEA-AD | not_assessed | All non-neuronal cells (glial and non-neural types) | |
| 5 | Lamp5 Lhx6 | subclass | SEA-AD | not_assessed | Lamp5_Lhx6 interneurons | |
| 6 | Lamp5 | subclass | SEA-AD | not_assessed | Lamp CGE interneurons | |
| 7 | Pax6 | subclass | SEA-AD | not_assessed | Pax6 CGE interneurons | |
| 8 | Sncg | subclass | SEA-AD | not_assessed | Sncg CGE interneurons | |
| 9 | Vip | subclass | SEA-AD | not_assessed | Vip CGE interneurons | |
| 10 | Sst Chodl | subclass | SEA-AD | not_assessed | Sst Chodl projecting MGE interneurons | |
| 11 | Sst | subclass | SEA-AD | down | Sst MGE interneurons | |

User-provided files must be either csv or gzipped csv files and have the same restrictions for file locations as described above. There are minimal restrictions on what can be included in this table:

- **cell_type:** This column includes the name of any (and ideally all) metadata items. ACE uses this column for two purposes: (1) to order all metadata values and (2) to find additional information about a metadata item to display.
- **direction** (optional): this field is currently the only way for coloring metadata on the "Explore individual annotations" tab (more details below). This tab looks for a column called "direction" which can be "down", "up", "unchanged", or "not assessed" (anything else is ignored) to determine color-coding and labeling.
- **[any other columns]**: there are no restrictions on any other column names. Information about whatever other columns are included in this table will be reported when the metadata value is selected.

Uses of the metadata table currently include:

1. **Ordering of cell types for confusion matrices and river plots**: Categorical annotations are ordered (e.g., sorted) in one of three ways. If provided as factors (e.g., in feather or h5ad files, or with extra columns with numbered "_id" and "_color", which we don't recommend), factor levels are used in ordering. Otherwise, the order that an annotation appears in the metadata file is the assumed order of cell types in the horizonal axis of the confusion matrix and the first column of the river plot (with missing values added at the end). Any variable where neither option is available is sorted alphabetically. Other annotations are reordered to maximize the diagonal on the confusion matrix and to minimize overlap in river plot "rivers".
2. **Display and color-coding on the "Explore individual annotations" tab**: Information from this table about each cell type annotation (e.g., individual table rows) are displayed when the relevant cell type is selected on the "Explore individual annotations" tab. Also, if a "direction" column is provided in the metadata table, plots on this panel can be color-coded by the directionality of a cell type's change with disease (e.g., AD). *If you have ideas on a better approach to color-coding, please provide feedback!*

Efforts linking ACE with existing and in process tools for cell type annotation are underway. This guide will be updated as integration progresses.

-----------------------------

#### ‘Upload or select data set’ pane

This pane allows you to choose preset annotation tables to compare or to point to your own tables and is the starting point for all other ACE functionality.

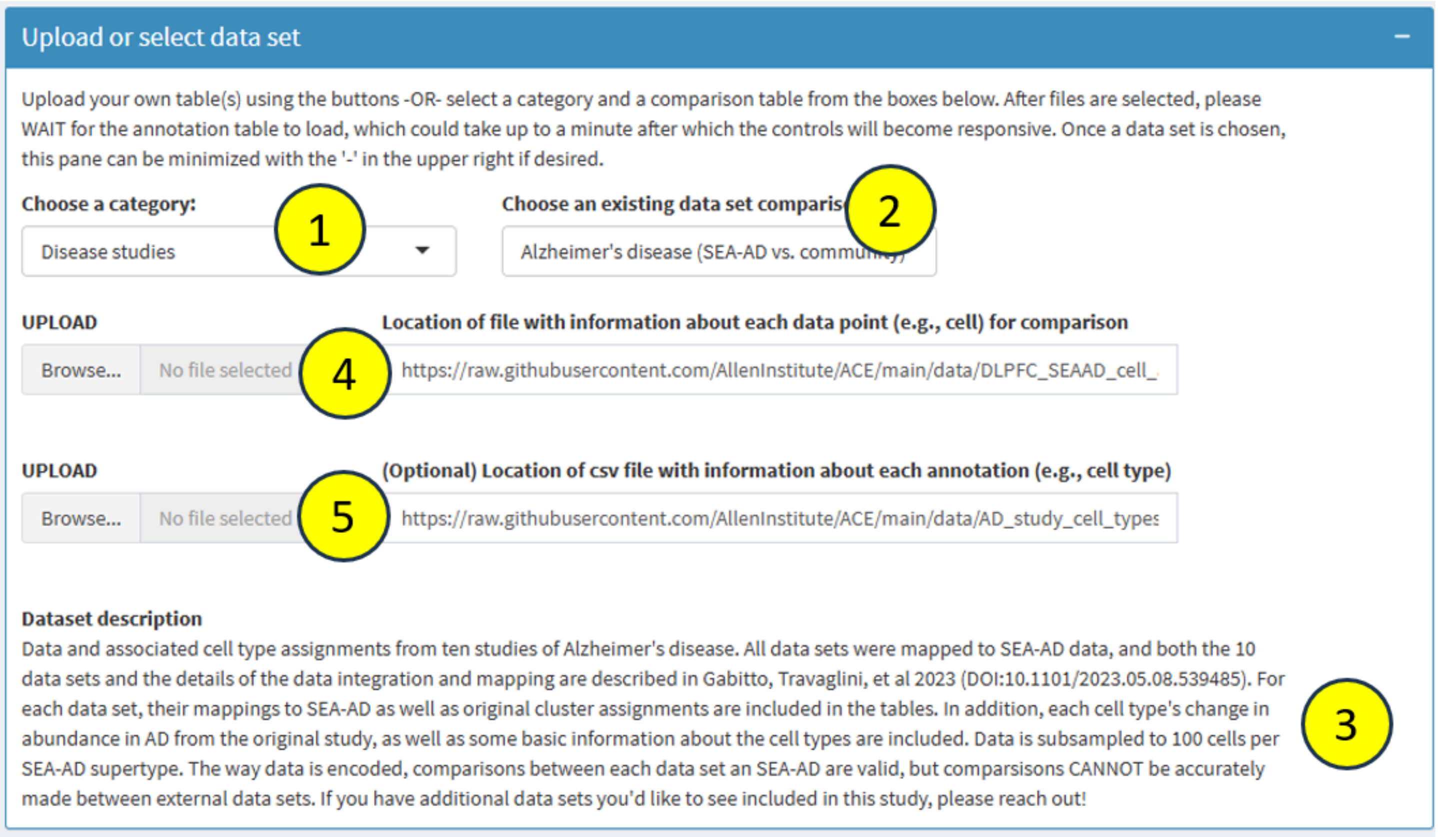


This pane includes a few components:

1) **Choose a category**: Choose from one of a few categories of pre-specified annotation categories (described above) or select “Upload or enter your own location” to point to your own data and then skip to #4.
2) **Choose an existing data set comparison**: Choose from one of the pre-specified annotation tables associated with this category. When you select a table, a few things will happen immediately:
    a. File locations will populate in #4 and #5. Do not change these file locations.
    b. The data will start to load. Depending on the table, this could take a few seconds up to a minute.
    c. Text will appear in the “Dataset description” section (#3). Read this text.
3) **Dataset description**: When a comparison table is selected, the text in this box will update to describe the content of the selected annotation table. After you read this text you can ignore #4 and #5.
4) **Location of file with information about each data point (e.g., cell) for comparison**: Either enter the location of the primary table here OR click on the UPLOAD button to upload a csv file. If an invalid file location is entered, you will see this error to the right of the bar “ENTER VALID CELL ANNOTATION FILE” and ACE will not work.
5) **(Optional) Location of csv file with information about each annotation (e.g., cell type):** Either enter the location of the metadata table here OR click on the UPLOAD button to upload a csv file. If you leave this blank or enter an invalid location, this table is ignored.

At this point your data set should be loaded, and you are ready to start filtering your data!

-----------------------------

### 'Filter cells in a dataset' pane

Cell filtering is a critical component of ACE, as it allows you to define the context for all the visualizations and statistics. For example, since there are >5000 cell types in mouse whole brain, it is not practical to perform comparisons on all of them at once (and some visualizations may not work properly with that much data). What is practical, and much more useful, is to see how all the inhibitory cells collected from mouse visual cortex map between cell type taxonomy versions, which dramatically decreases the size and scope of the data sets.

**Example: Filtering by categorical variables**

As another example (shown below), we can restrict our visualizations to only non-neuronal cells collected from a previous study of AD that successfully map to SEA-AD cell type. Here is how that filtering looks:

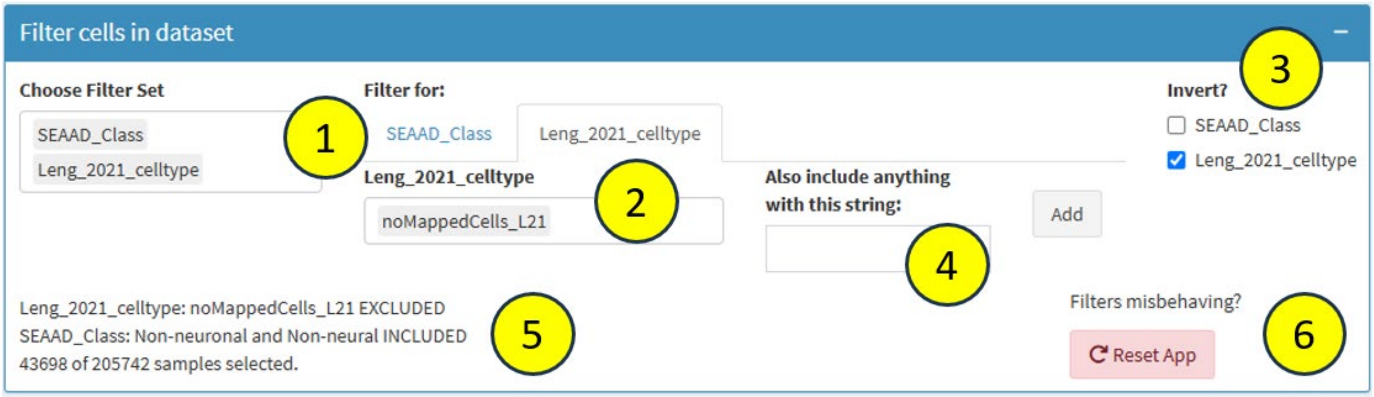


The "Choose Filter Set" box (#1) allows you to select one or more metadata columns on which to perform the filtering. This box (and all other boxes in the app where you can type) have autocomplete and will help you select possible options to select. For each chosen variable, the "Filter for" box (#2) allows you to choose how to filter the data. Categorical variables can be filtered by listing all of the values for that variable that you'd like to include in the visualizations for the app. For example, to retain all non-neuronal cells, you could either set the "SEAAD_Class" filtered to "Non-neuronal and Non-neural" (as done in this example) or you could set the "SEAAD_supertype" filter to include every non-neuronal cell type (or both). Categorical filters can also be "inverted", meaning that all cells except the ones filtered will be included (#3). In this case, all cells that failed to map to SEA-AD (and therefore have a "Leng_2021_celltype" value of "noMappedCells_L21") are excluded.

Another filter (that is not used in this example) is the "Also include anything with this string:" filter (#4) which will add any metadata with a particular string to the existing set of metadata. For example, if we wanted to include all cell types with the string "SFG", we could set our filters as follows.

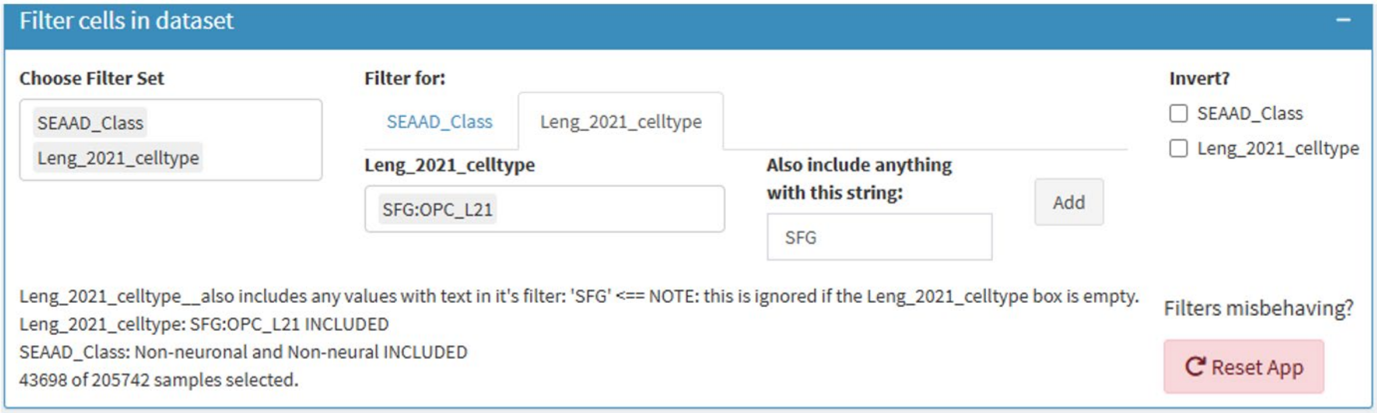

Finally, the bottom left corner of this box filtering box (#5) shows the filters that have been applied, whether cells from that filter and included or excluded, as well as the number of cells out of the total table that will be included for other app components. The two filtering options above would result in an identical set of cells selected because all cell types in the "Leng 2021" study start with the characters "SFG".

*We note that the "Invert?" and the "Also include anything with this string:" filters can occasionally get stuck in the wrong state. If this happens, we recommend refreshing the app (#6) to re-select or re-upload your data set.*

**Example: Filtering by numeric variables**

In this example, we can restrict our selection to only include data points that fall within a specific numeric range. Here we are looking at patch-seq cells from mouse visual cortex that correspond to GABAergic interneurons ("Mouse visual cortex (GABAergic neurons)"). The following configuration would set ACE to only consider cells with "Resting membrane potential (mV)" between -80 and -60:

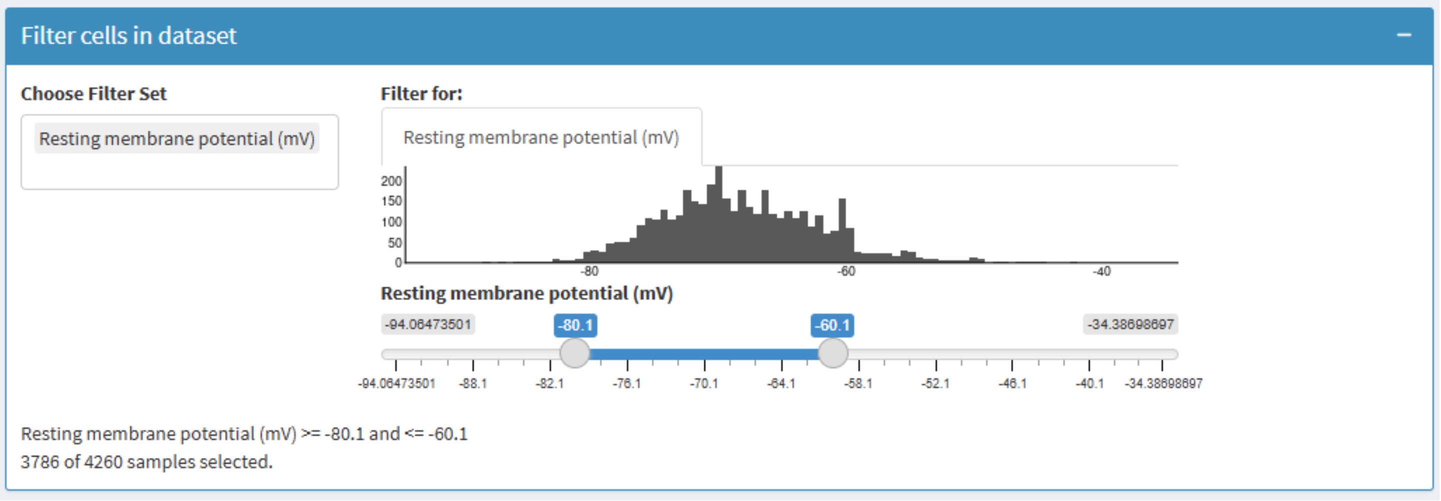


In this case we see a slider bar which allows you to select the range of values to include (shown in blue beneath) as well as a histogram showing how many data points include values at a specific range (above the slider). In this case we see that the most common resting membrane potential is ~70mV and that most cells fall in a range thereabouts. Other components of the filter look and work the same as for categorical variables.

----------------------------

### Visualizations and statistics: Intro

Once the annotation table and relevant rows are chosen, all functionality of ACE takes place in the 'Visualizations and statistics' pane. These tabs provide different tools for comparing cell-level annotations and (optionally) providing additional information about each annotation. The "Intro" tab provides a brief overview of the other visualization tabs, with more details described in the user guide sections below.

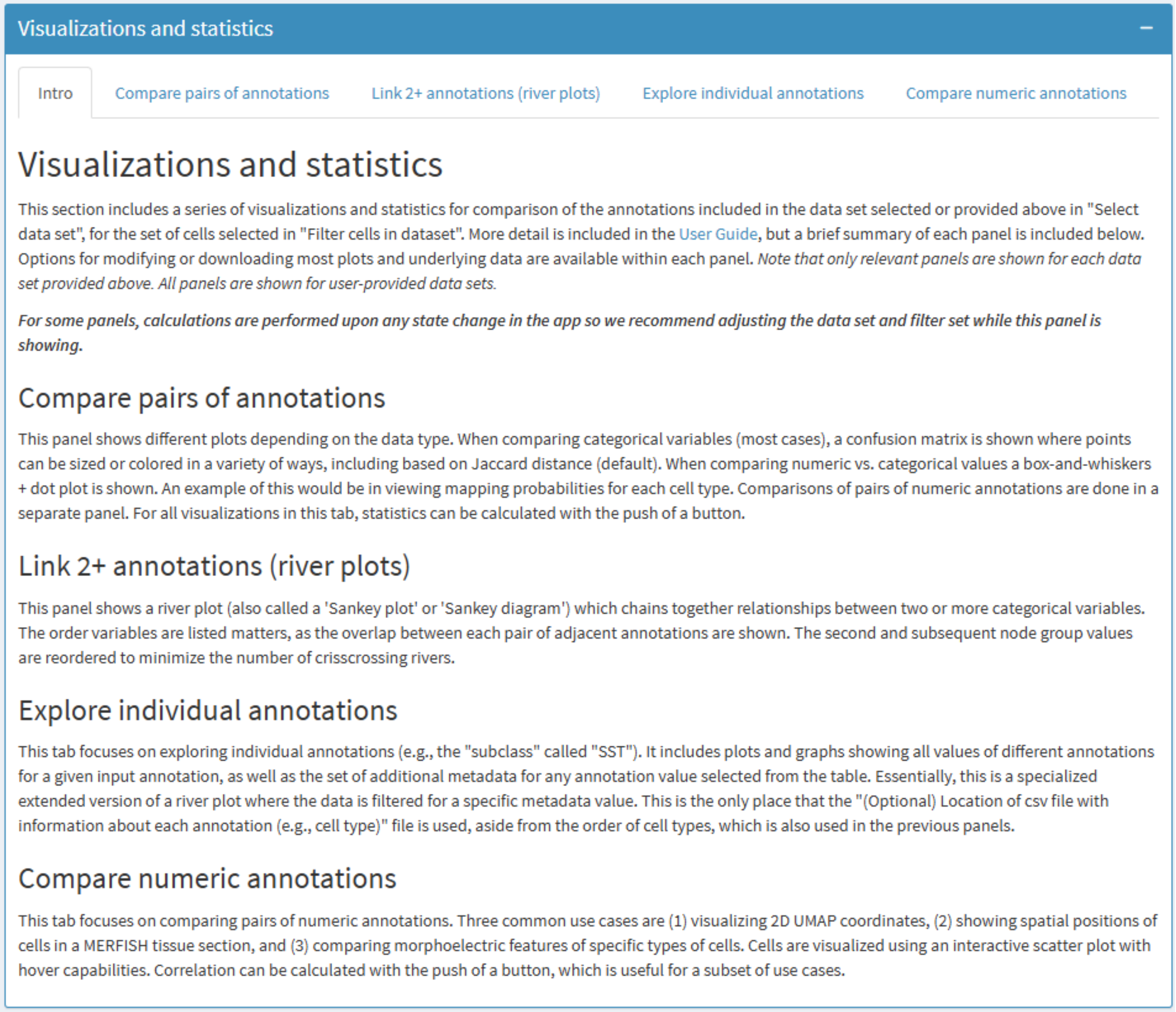
Visualizations and statistics

Intro | Compare pairs of annotations | Link 2+ annotations (river plots) | Explore individual annotations | Compare numeric annotations

#### Visualizations and statistics

This section includes a series of visualizations and statistics for comparison of the annotations included in the data set selected or provided above in "Select data set", for the set of cells selected in "Filter cells in dataset". More detail is included in the User Guide, but a brief summary of each panel is included below. Options for modifying or downloading most plots and underlying data are available within each panel. *Note that only relevant panels are shown for each data set provided above. All panels are shown for user-provided data sets.*

***For some panels, calculations are performed upon any state change in the app so we recommend adjusting the data set and filter set while this panel is showing.***

##### Compare pairs of annotations

This panel shows different plots depending on the data type. When comparing categorical variables (most cases), a confusion matrix is shown where points can be sized or colored in a variety of ways, including based on Jaccard distance (default). When comparing numeric vs. categorical values a box-and-whiskers + dot plot is shown. An example of this would be in viewing mapping probabilities for each cell type. Comparisons of pairs of numeric annotations are done in a separate panel. For all visualizations in this tab, statistics can be calculated with the push of a button.

##### Link 2+ annotations (river plots)

This panel shows a river plot (also called a 'Sankey plot' or 'Sankey diagram') which chains together relationships between two or more categorical variables. The order variables are listed matters, as the overlap between each pair of adjacent annotations are shown. The second and subsequent node group values are reordered to minimize the number of crisscrossing rivers.

##### Explore individual annotations

This tab focuses on exploring individual annotations (e.g., the "subclass" called "SST"). It includes plots and graphs showing all values of different annotations for a given input annotation, as well as the set of additional metadata for any annotation value selected from the table. Essentially, this is a specialized extended version of a river plot where the data is filtered for a specific metadata value. This is the only place that the "(Optional) Location of csv file with information about each annotation (e.g., cell type)" file is used, aside from the order of cell types, which is also used in the previous panels.

##### Compare numeric annotations

This tab focuses on comparing pairs of numeric annotations. Three common use cases are (1) visualizing 2D UMAP coordinates, (2) showing spatial positions of cells in a MERFISH tissue section, and (3) comparing morphoelectric features of specific types of cells. Cells are visualized using an interactive scatter plot with hover capabilities. Correlation can be calculated with the push of a button, which is useful for a subset of use cases.

Each remaining tab within this pane corresponds to a different visualization that applies to the filtered data set defined above. **Note that not all functionalities will be available for every annotation table**. Depending on the types of data provided in the input tool, certain tabs may not work or may be omitted. Nonfunctional tabs will be hidden for all annotation tables listed on ACE, whereas all tabs are always shown for user-provided tables whether they are functional or not.

Under the hood, the "Intro" tab provides a calculation-free home page for this pane. Since some calculations are performed upon any state change in the app, we recommend adjusting the data set and filter set while this panel is showing, rather than while one of the main visualization tabs are shown.

----------------------------

### Visualizations and statistics: Compare pairs of annotations

This tab allows comparison of any two pieces of categorical or numeric metadata. All of the tabs are laid out roughly the same way, with controls at the top, the main visualization in the middle, and download or other content on the bottom, details of the controls for this tab are shown a bit below.

#### COMPARING TWO CATEGORICAL VALUES

An example of the 'Compare pairs of annotations' tab is shown below, with default parameters for comparing a pair of categorical variables:

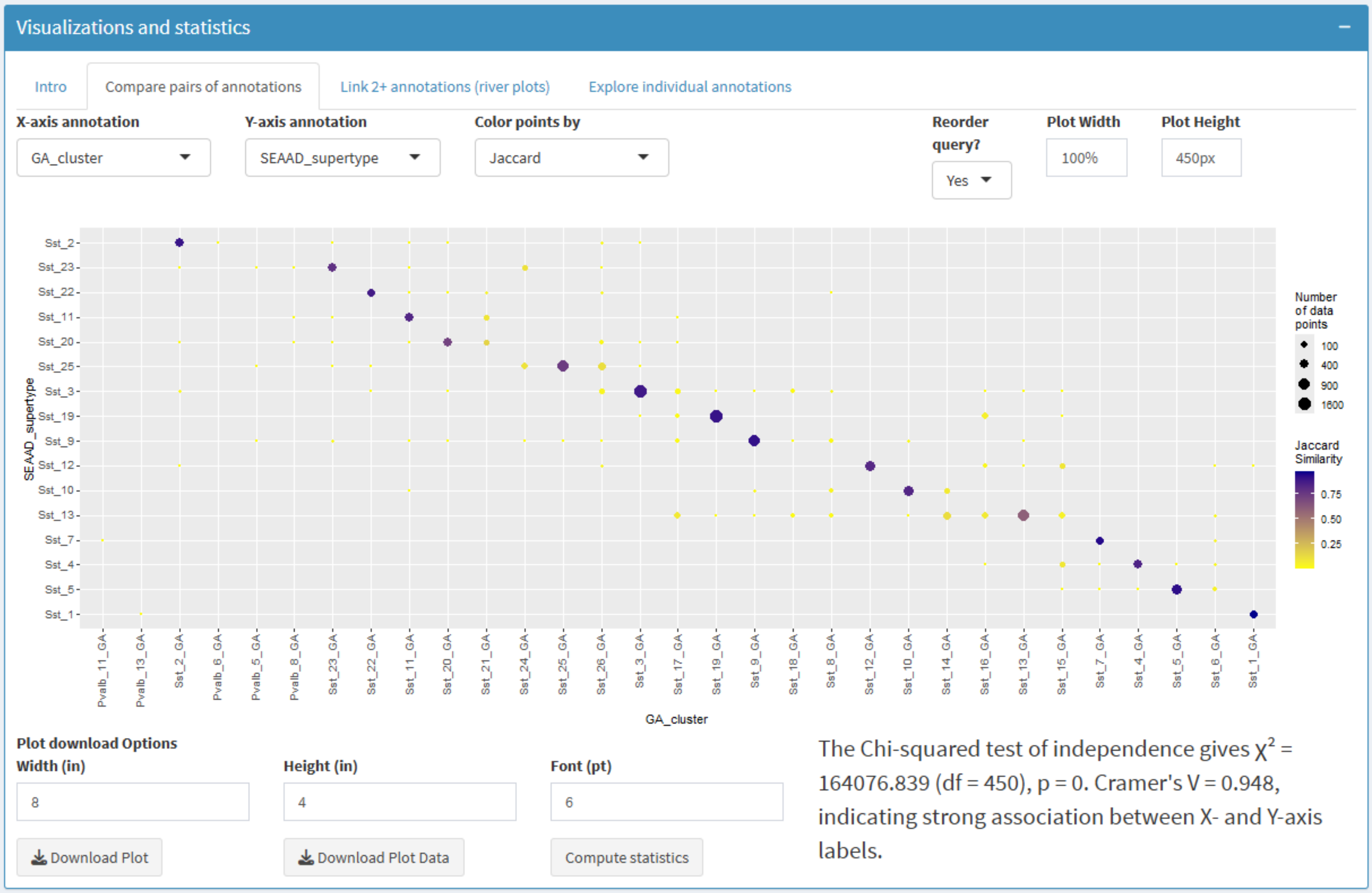


This example compares cell type assignments of neurons defined as SST in SEA-AD with their original cell type assignments in the great ape (GA) study. The confusion matrix shows a high correspondence between cell type definitions in these two studies, with most cells from SEA-AD supertypes (Y-axis) assigned to 1-2 clusters in the great ape study (X-axis), and strong statistical confirmation of significance. This result makes sense, since SEA-AD supertypes were defined using great ape assignments as a starting point. By default, Jaccard distance is shown, with blue values indicating a better correspondence and yellow values indicating a poorer correspondence. We note that values on the X-axis are ordered to match their appearance in the metadata annotation file (in this case representing the order of cell types in the GA taxonomy).

We can also change the plot to show the proportion of cells of each SEA-AD supertype assigned to each GA clusters (e.g., rows sum to 1):

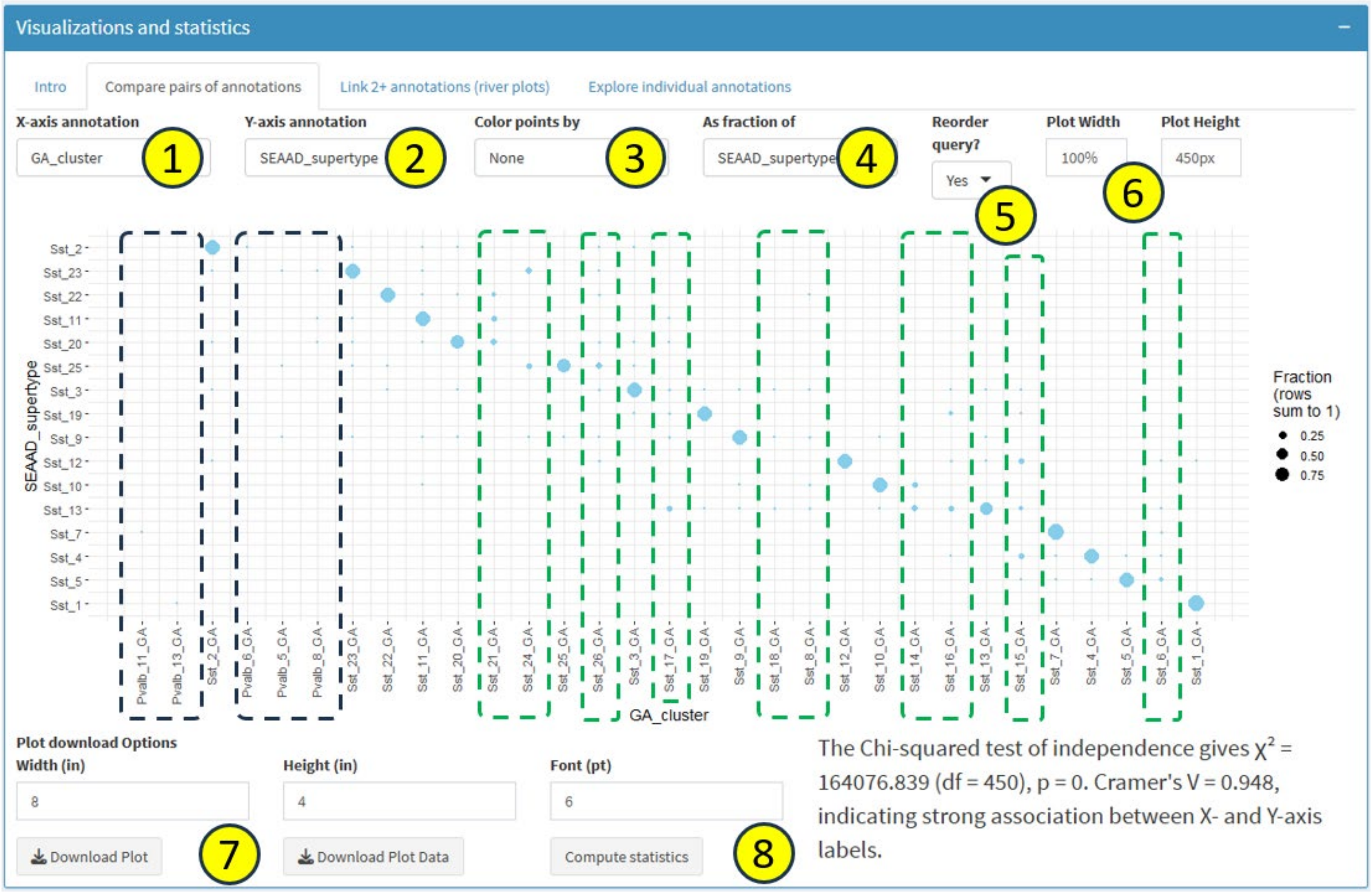


Again, we see that nearly all of the circles are either very big or tiny/missing, indicating good agreement in annotations. We also see a very small number of cells (tiny dots) that were defined as PVALB in GA but are not reassigned to SST types (black hashed boxes). Since the number of cells is small, and these subclasses are not perfectly distinct, such an overlap is nothing to be concerned with. Finally, we note that matching tables in both taxonomies (e.g., Sst_1 vs. Sst_1_GA) line up because of how these cell type taxonomies were made—this is typically NOT the case. Several clusters were removed between GA and SEA-AD taxonomies (green hashed boxes) with cells being reassigned new SEA-AD cell type assignments. *The green and black hashed boxes are added for demonstration purposes and are not part of the ACE GUI.*

This tab includes the following controls, which can vary slightly depending on selections:

1) **X-axis annotation**: which annotation will be shown on the horizontal axis?
2) **Y- axis annotation**: which annotation will be shown on the vertical axis?
3) **Color points by**: Which metadata should points be colored by? For categorical visualizations the default value is Jaccard, which produces a plot like the one two figure above. In this case, color represents Jaccard similarity (bigger values and bluer colors represent higher similarity), while the size of the point is scaled by the number of cells in each intersection. To change this scaling (using "As a fraction of" below), first change the "Color points by" drop down to any of the other values.
4) **As a fraction of**: This drop-down is only shown if "Color points by" is not set to "Jaccard". Otherwise, there are three options: "None" (default), in which case the size of the points corresponds to the number of cells in that intersection. If changed to the metadata shown in y-axis or x-axis annotation, this converts the points to fractions where the values row- or column- sum to 100%. This is one way to customize the relative size of each point; however, it will not change the absolute size of the points (see below).
5) **Reorder query?**: This parameter will attempt to sort the entries within the y-axis metadata to create as good looking of a diagonal as possible. This is extremely useful if the metadata are not sorted in the same way (or at all) and is set to 'Yes' by default. If set to 'No', the original order is retained.
6) **Plot Width / Plot Height**: Parameters that will change the width (in %) and height (in pixels) of the visualization on your screen. Note that you need to include the "%" sign in the Plot Width box.

ACE also includes some functionality for **downloads and statistical analysis** in several panes:

7) **Download Plot / Download Plot Data:** The bottom of this tab (and several others) includes an option to download the current visualization, along with a few parameters for the size of the images and included text. These defaults are not always reasonable, so it may be worth trying a few downloads to see which one looks best. The "Download Plot" button defaults to using a generic file name and file location, so we recommend changing the names as appropriate. Beside this button is a "Download Plot Data" button, which will download the relevant numbers shown for the plot on the page. In the case of confusion matrices, rows and columns will correspond to the two selected annotations and matrix values will correspond to the counts of data points with that pair of annotations.
8) **Compute statistics**: When this button is clicked a statistical test will be performed related to the plot currently shown on the screen. For pairs of categorical variables (like those shown above), a Chi-Squared test is run to assess whether the confusion matrix shown is non-random, and the Cramer's V test is run to determine how strong the relationship is (on a scale from 0 to 1).

**Customizing point size:** It is possible to customize the point size in a couple of ways. The relative size of points can be adjusted using the "Color points by" and "As a fraction of" controls above. Second, when downloading the image using the "Download Plot" button in the bottom left, adjustment of the width and height will impact the absolute size of the points (larger images = smaller points), while retaining the relative sizes from the GUI.

#### COMPARING NUMERIC WITH CATEGORICAL VALUES

In addition to comparing categorical variables, ACE can compare categorical and numeric variables. In this case, a "swarm" plot is shown, where each point represents a single cell, the width of the cells at a given vertical position showing the relative density, and an overlaid mean +/- interquartile range shown as red and black ticks, respectively. A color legend listing all (up to 25) color annotations is shown on the right. Here is an example:

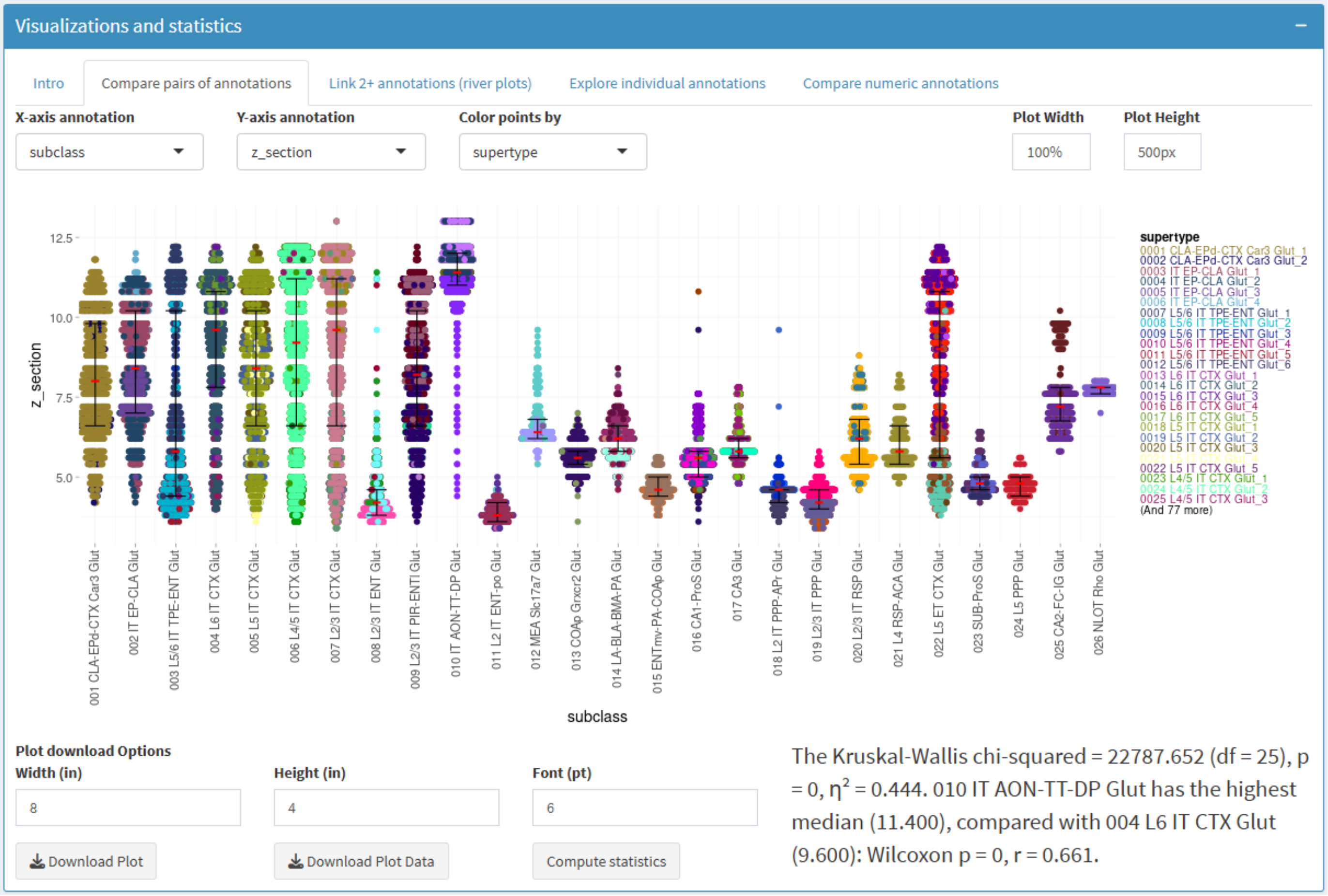

In the plot above we use the example of mouse MERFISH data, showing the z-section from which all class="01 IT-ET Glut" cells are located, divided by subclass and color-coded by supertype. We see that some subclasses tend to be more rostral (low values) than caudal (high values), while others have different supertypes spanning the brain (e.g., subclass 022 has teal-colored cells rostrally and purple-color cells caudally). These types of plots are also useful for assessing data quality by reviewing the spread of a QC metric across cells of a given annotation (e.g., what fraction of SST cells have high mapping, do cells from a single donor have more mitochondrial reads than for others), among other purposes.

These plots include the same controls and buttons as for comparison of categorical variables. In this case the "Download Plot Data" button will download a 3 x N matrix, where the rows correspond to each data point and the columns correspond to the sample_id, x-axis annotation, and y-axis annotation, respectively. In this case the statistics return to a Kruskal-Wallis test indicating the difference from chance in the distribution across all categories along with a Wilcoxon test comparing the two categories with the highest values.

----------------------------

#### Visualizations and statistics: Link 2+ annotations (river plot)

This panel creates an interactive river plot (also known as a "Sankey diagram") that allows you to compare two or more pieces of metadata (see below). In the Node Groups box (#1) you can enter any number of metadata fields in order (the order matters!) to generate the river plot. For each metadata field you get a stacked bar plot showing the number of cells that have each value within that metadata, with the value for each label shown (#2). In addition, for each adjacent pair of bar plots, you get "rivers" connecting each pair of metadata values together, where the thickness of each line represents the number of cells sharing the corresponding values from the two metadata fields (#3). Since this plot is interactive, you can hover over any bar or river and see what values and numbers correspond to what you are hovering over (#4). On the right side of the plot (#5) there are extra controls that allows you to zoom, pan, and interact with the plot in various other ways. Finally, you can download the plot using the buttons on the bottom (#6; as described above). No data downloads or statistics are available for river plots.

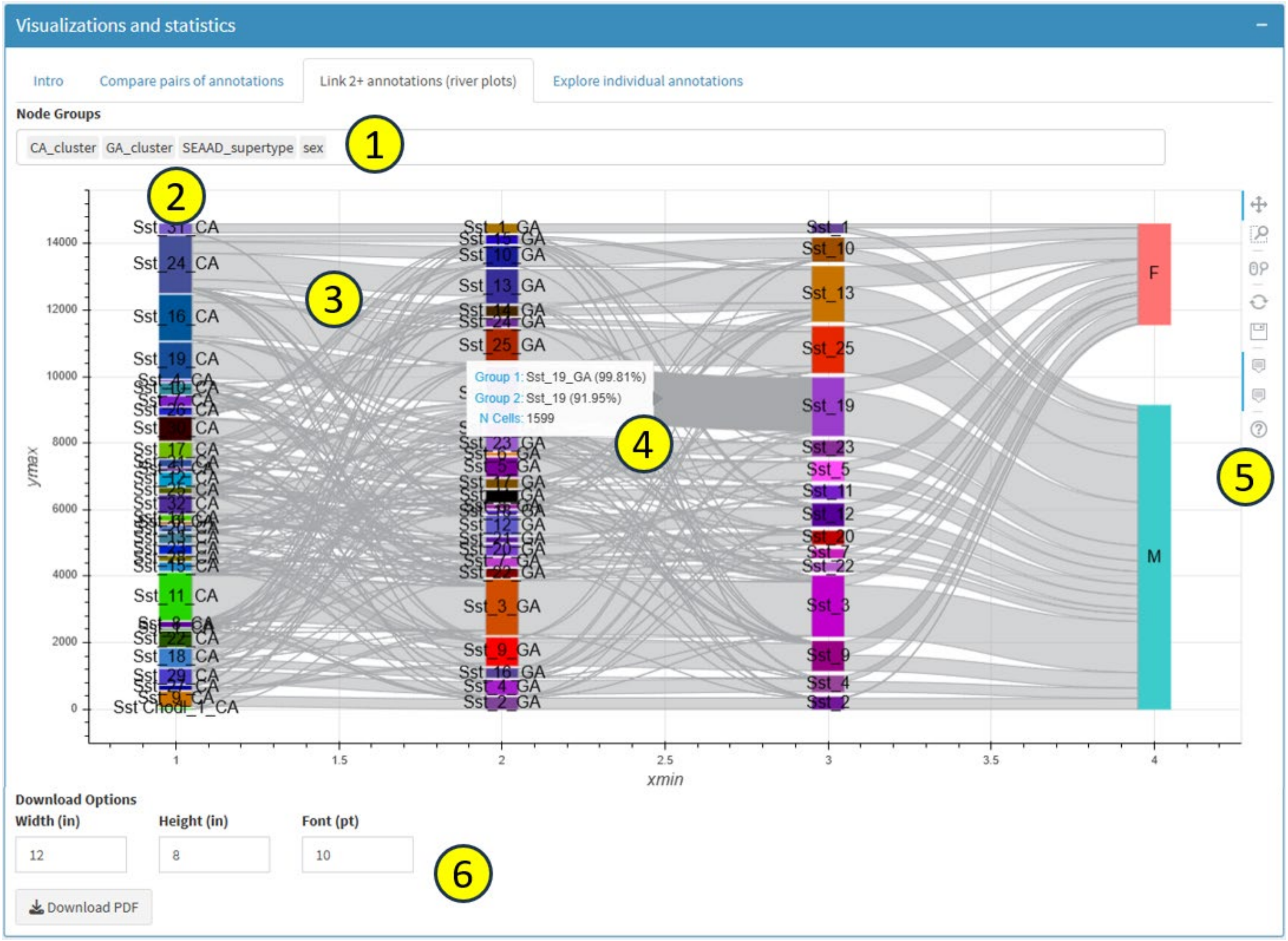


The order of the leftmost column is pre-defined (either by factor levels, by the inputted metadata table, or alphabetically). Entries in the remaining columns are sequentially reorders in an attempt to minimize the crisscrossing rivers and maximize the horizontal flow between annotations (Node Groups).

Currently there is no way to resize the box or to show the same metadata more than once in the Node Groups sequence, and this may or may not be updated later. *Do not resize the screen while river plots are active!*

-----------------------------

### Visualizations and statistics: Explore individual annotations

As the name suggests this tab is focused on exploring individual annotations (see below). *In essence, this tab shows basically the same information as river plots for a single metadata value, but without showing the rivers and with extra information about metadata fields optionally shown.* This can be useful for manual annotations of individual clusters (or supertypes, subclasses, etc.) or for understanding how cell types defined in one study compare with cell types defined in other studies. This is the only tab that uses information from the metadata information table beyond the ordering of the annotations. In this case we use the "Alzheimer's disease (SEA-AD vs. community)" data set as an example.

**(Optional) Location of csv file with information about each annotation (e.g., cell type)**

https://raw.githubusercontent.com/AllenInstitute/ACE/main/data/AD_study_cell_types_for_app.csv

Unlike the other tabs which compare all values for two or more pieces of metadata, this tab is anchored around a single "**Annotation value**" from a single "**Annotation group**" or metadata column (#1 below), which will show up in the left-most position in the visualizations on this tab. Once this is chosen, the rest of the tab will be performed in comparison to that one annotation value. We next indicate which other "**Comparison groups**" (e.g., specific metadata columns) will be compared against the starting annotation value (#2 below). In this case, we choose mappings from the eight other AD studies that include neuronal types:

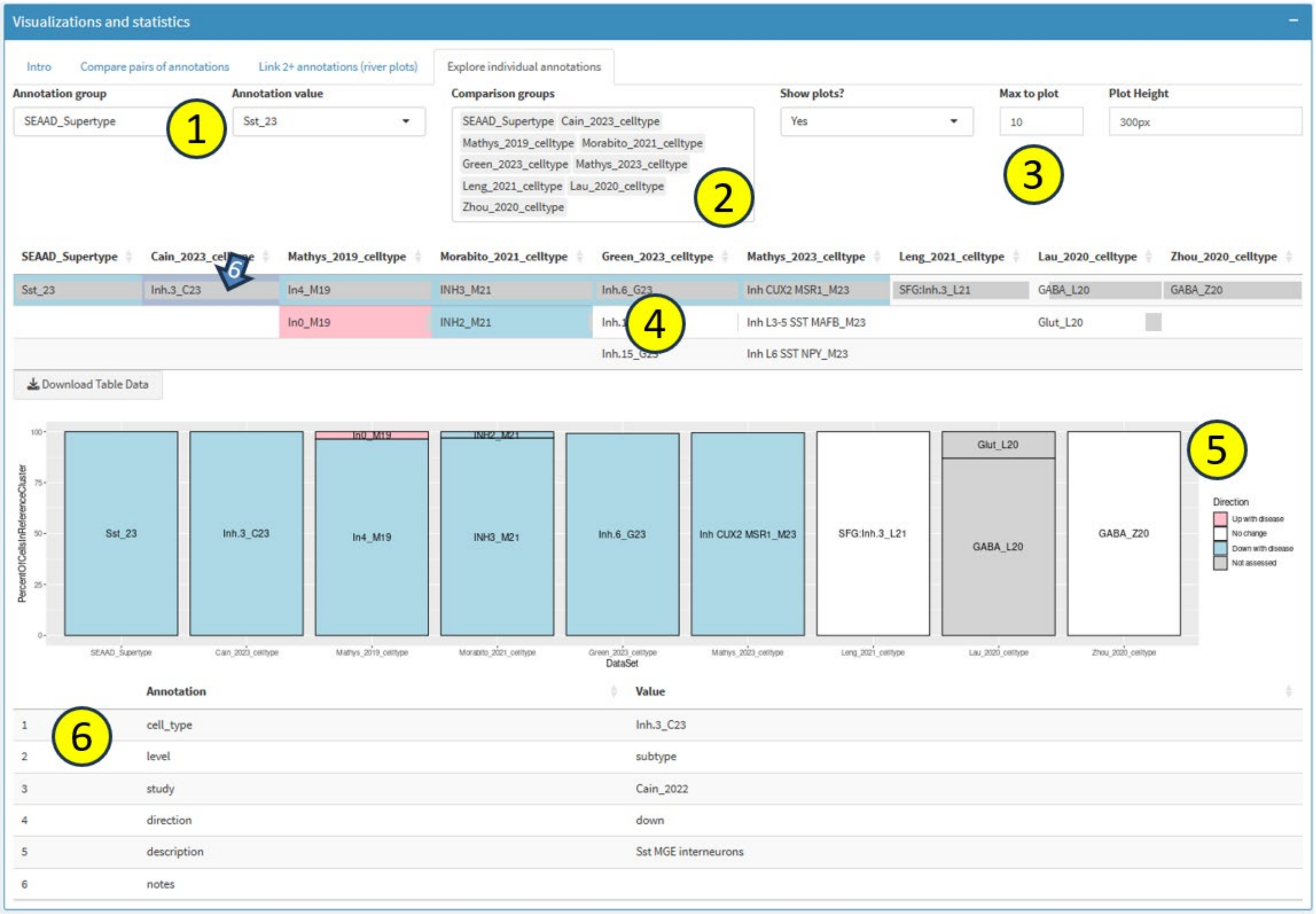

The other controls at the top describe whether/how comparison information will be displayed as a bar plot below the chart display (#3, more on that shortly). Specifically:

- **Show plots?:** A Yes/No call on whether the bar plot should be shown
- **Max to plot:** The maximum number of top overlaps to show in the bar plot (for example, if there are 20 SST clusters and "Annotation value" is set to SST subclass, then only the 10 most abundant SST clusters would be shown in the plot).
- **Plot Height**: Height of the plot in pixels.

The next section (#4) shows a chart of the selected annotation data along with the corresponding values from the comparison groups, sorted descending from highest to lowest overlap. Within each table entry, a grey bar indicates the fraction of cells from the selected annotation value that also have the comparison value listed. Boxes are also color-coded with whether they values are up (red), down (blue), or unchanged (white) with disease. These calls rely on reading the "direction" column values in metadata information table. For both the chart and bar plot views, no color-coding is done if the "direction" column is omitted. A "Download Table Data" button is included for downloading the numbers behind the grey boxes (and the "direction" values, if provided) for annotations shown in the table (and in the bar plot, if shown). This download includes all entries, not just the top "Max to plot".

If shown, the bar plot (#5) displays the same information as the chart using a different graphic, with each bar indicating the fraction of cells from the selected annotation value that also have the comparison value listed, again colored by change in AD. The colors and order of information in the two graphics should match, except that a distinction is made between annotations which do not increase or decrease in disease because there really isn't a change ("No change") or because no comparison was performed ("Not assessed"). If direction data is not provided, these are color-coded a different shade of grey and given the label "Not provided".

The final part of this tab is a table that displays additional provided information for a given metadata value (#6). If you click on any box in the chart described in #4 (example above), then a table showing the provided metadata for that specific cluster will be shown below. We anticipate that future updates will allow this information to be collected from standard cell type taxonomy formats in addition to user-provided csv files.

------------------------------

### Visualizations and statistics: Compare numeric annotations

This tab focuses on comparing pairs of numeric annotations. Cells are visualized using an interactive scatter plot with hover capabilities. Three common use cases are (1) visualizing 2D UMAP coordinates, (2) showing spatial positions of cells in a MERFISH tissue section, and (3) comparison of QC metrics or cellular features (such as morphoelectric measurements). It's worth noting that there are already a number of interactive tools dedicated to 2D scatter plot visualizations in the context of single cell data, including the Allen Brain Cell Atlas, CZI CellXGene, Cirrocumulus, and Cytosplore, which may be more appropriate than ACE for some use cases.

This tab is only activated for annotation tables with at least two pieces of numeric metadata. Otherwise, the tab either will not be present in the Visualization and statistics pane (e.g., for prepopulated data sets), or will return a warning in place of a plot here (e.g., for user-provided data sets):

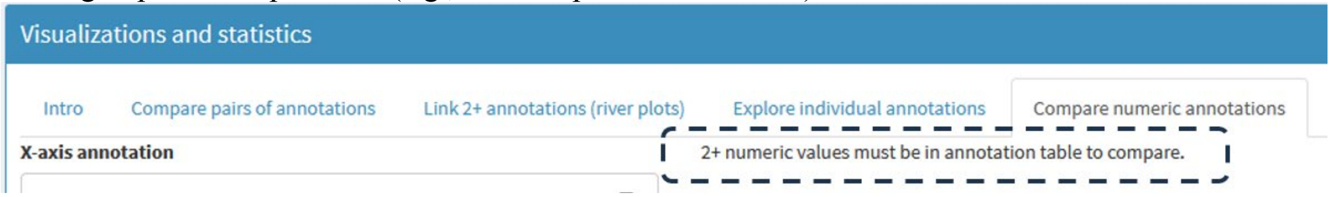


If two or more numeric metadata are available, a page like this will be displayed (in this case showing "Middle temporal gyrus (initial studies)":

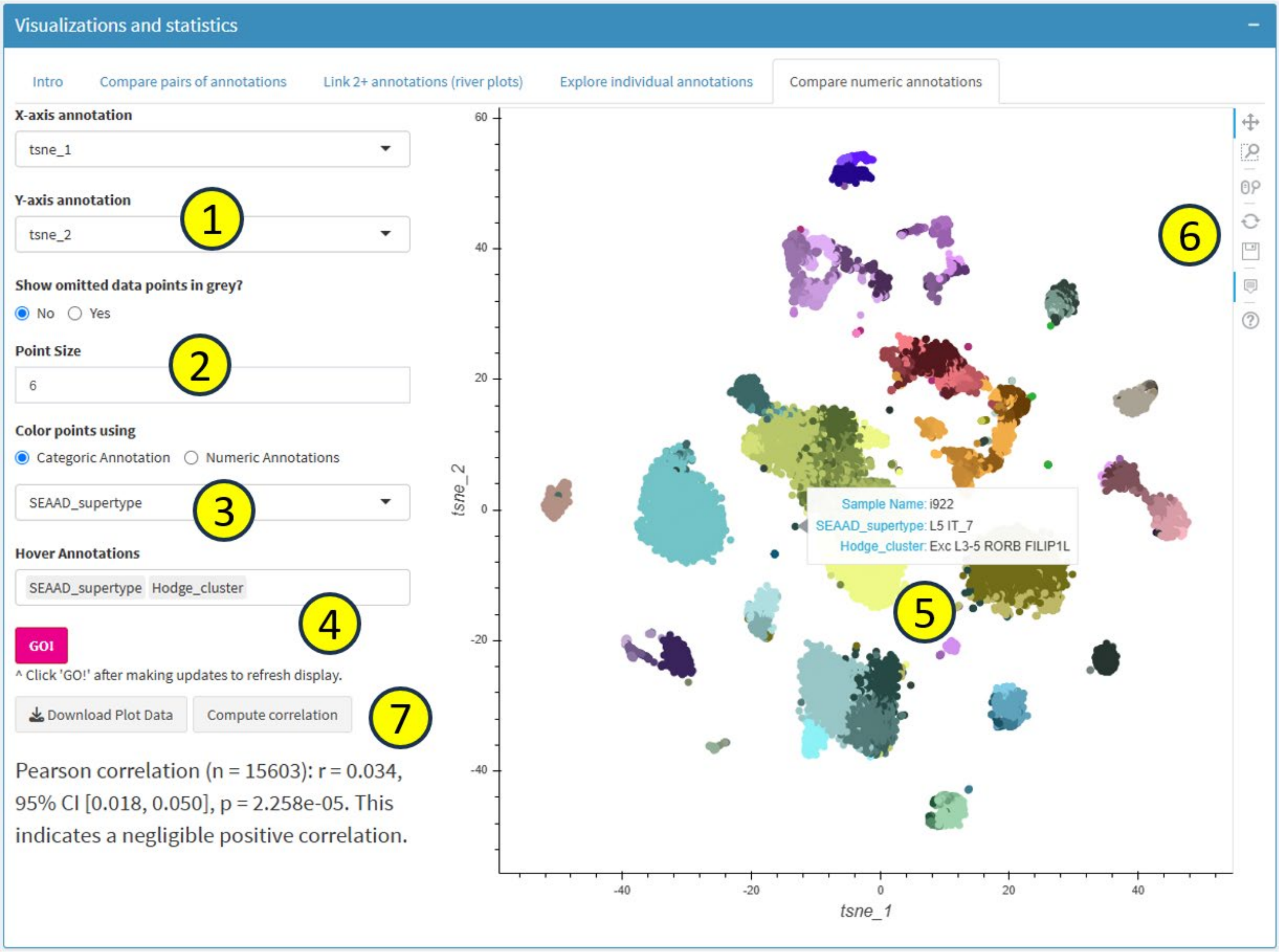

In the **X-axis annotation** and **Y-axis annotation** dropdowns (#1), you can select which annotation will be shown on the horizontal axis and vertical axes, respectively. For this pane only numeric annotations will be available to display, while categorical annotations used for other tabs will be hidden. You can also select which **Point Size** to use (#2), with the default (6) being reasonable in most cases.

The main use of this tool is for color-coding the points by metadata. You can color-code based on a single categoric annotation (#3 above) or based on up to three numeric annotations at once (alterative panel below).

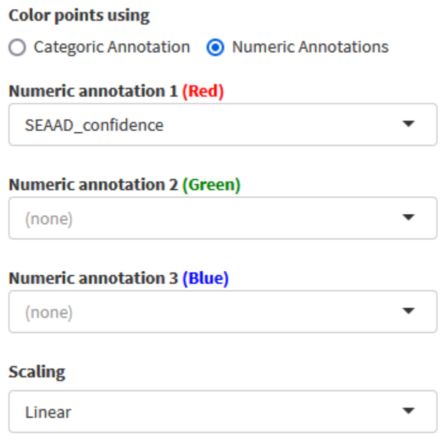


Since this plot is interactive, you can hover over any point (#5) and see specified information about that cell (#4). On the right side of the plot (#6) there are some extra controls that allows you to zoom, pan, and interact with the plot in various other ways. ***Note that to save your image, you need to use the icon in control panel on the right ( ). There is NOT a separate "Download Plot" button.*** However, as with other tabs, there are buttons to "Download Plot Data" and compute statistics (Pearson correlation) (#7). In this case, the Pearson correlation is quite low (r=0.034) because there is not expected linear relationship between tSNE coordinates of cells.

##### Omitting vs. deemphasizing omitted data

ACE provides two different ways to visualize scatterplot data when some cells are excluded (in this case, only Non-neuronal cells are shown). If "Show omitted data points in grey?" (arrow) is set to "Yes", all cells are included but colors are set to grey, and hovering is disabled in omitted cells. If set to "No" then these cells are omitted from the visualization entirely.

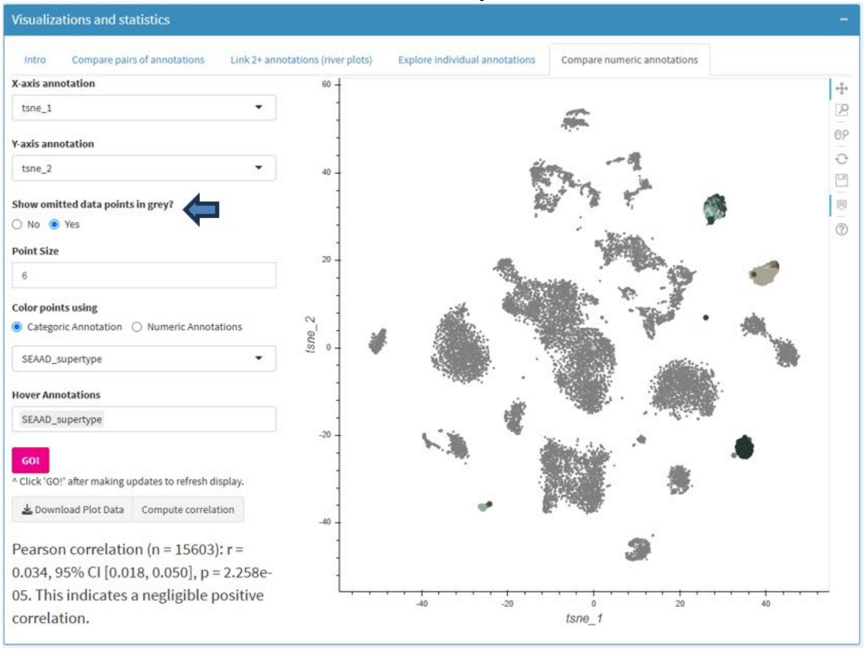


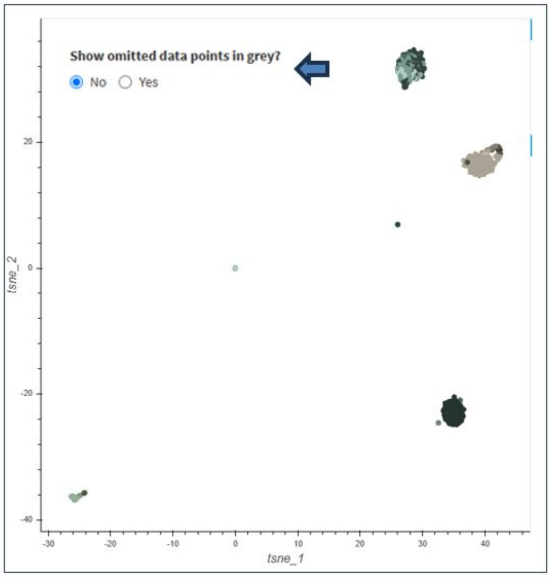

**Showing spatial positions of cells in a MERFISH tissue section**

The above example shows how to visualize metadata on a UMAP plot. Below we include an example of how to do the same on a single MERFISH tissue section. To view MERFISH cell coordinates, it is critical to first filter the data to only include cells from a single plane of section. In this case we use the filter pane to select a very narrow band in "z_section" corresponding to a single tissue section.

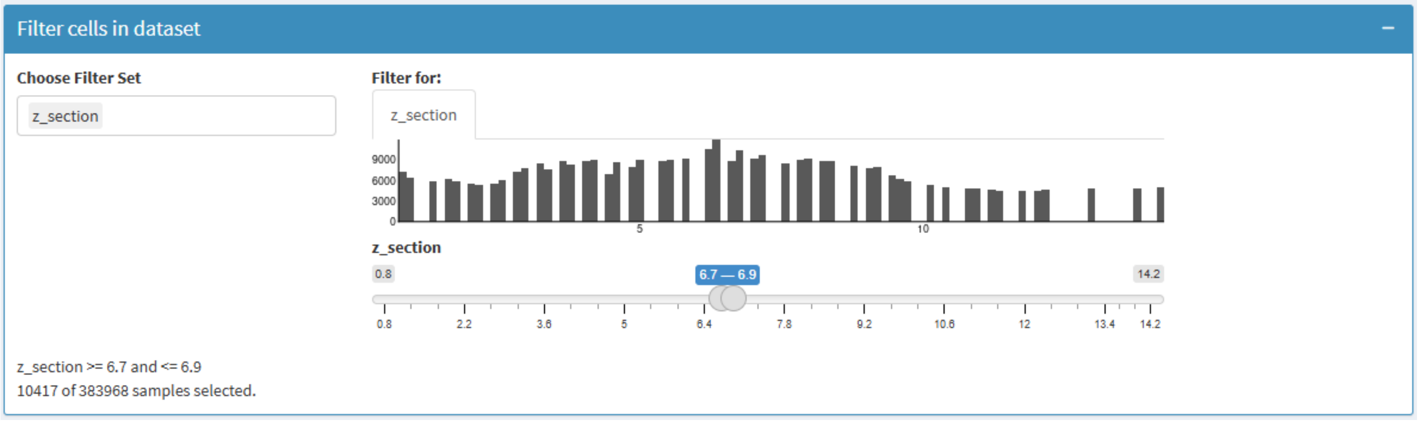


The image below then shows the subclass-level annotations for this section, with additional annotations available to view on hover.

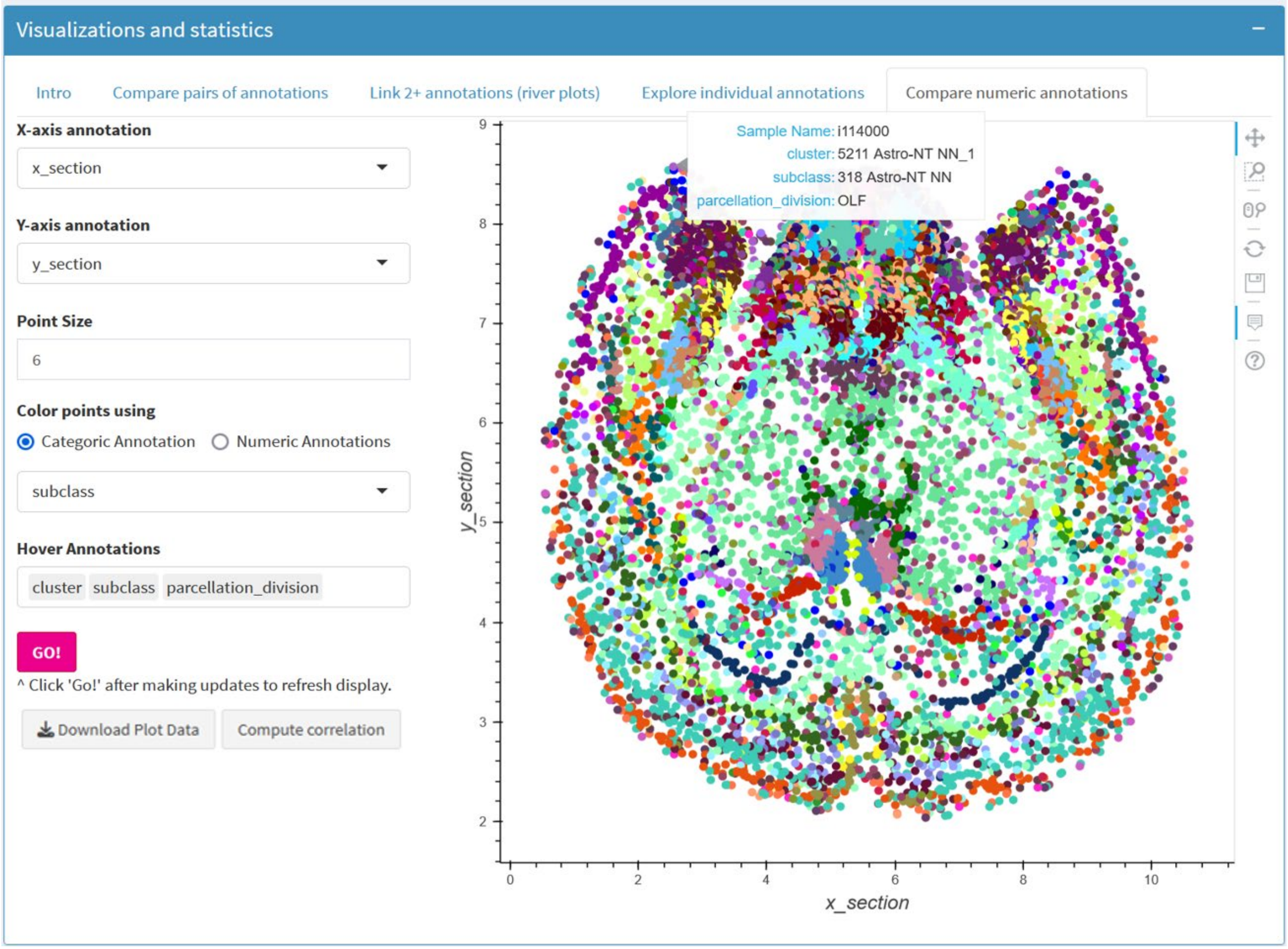


A future improvement to this tab is planned to allow a user to compare two separate pieces of metadata for the same cells (e.g., color code cells that are only in hippocampus, only inhibitory neurons, neither, or both).

----------------------------

### Use case: Comparing morphoelectric properties of different cell types

Electrophysiological differences between intratelencephalic (IT) and extratelencephalic (ET) cells are well established with ET neurons having a higher ratio of voltage at the peak and voltage at steady-state in response to hyperpolarizing current injections ("Sag ratio") than IT neurons in humans [13]. By exploring an available table of patch-seq data collected in mouse primary motor cortex and restricting the view to these two subclasses (ET and IT), this difference is clearly consistent between all ET and IT cell types, shows a statistical difference across cell types, matching previous reports in humans. Additional separation between mouse ET and IT cells can be seen when comparing Sag ratio (higher in ET) and action potential (AP) width (defined as the duration of the AP at half-amplitude from AP threshold; higher in IT) using a scatter plot. **So how do we show this?**

*Step 1: Select the data set*

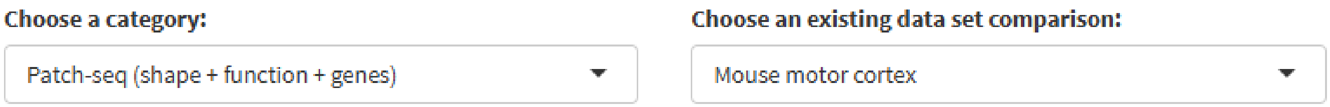


*Step 2: Filter data set to only include ET and IT cells*

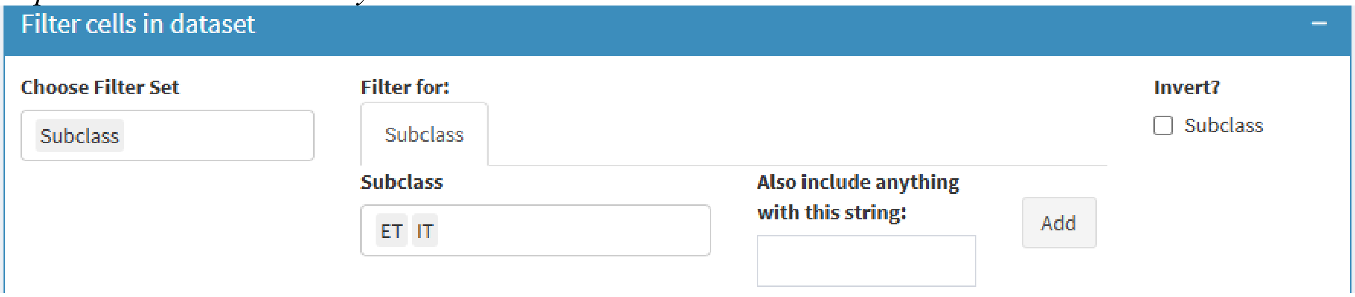


*Step 3: Plot Sag ratio vs. cell type*

Here we plot the Sag ratio vs. cell type ("Yao2021_CTKE_Ttype_M1") and color-code by Subclass. We then click the "Compute statistics" button to see that there is a statistical difference between cell types. We also find a statistically significant difference if we compare Sag ratio vs. subclass directly (not shown).

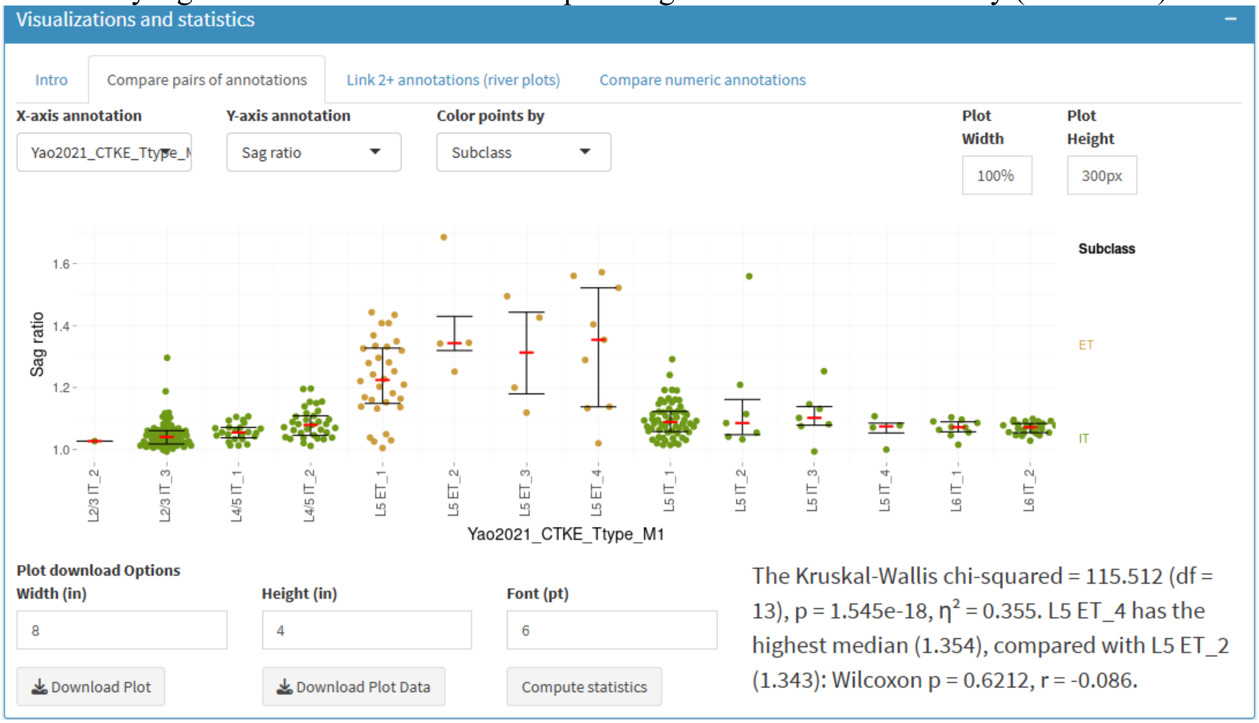

*Step 3: Plot Sag ratio vs. AP width*

Using the same filters, we next plot the Sag ratio vs. AP width (ms) and color-code by Subclass. We also add some hover annotations and click "Compute correlation". Altogether, this plot shows us that these two features alone do an excellent job of separating ET (yellow) from IT (green) cells, and furthermore, that the two metrics are negatively correlated.

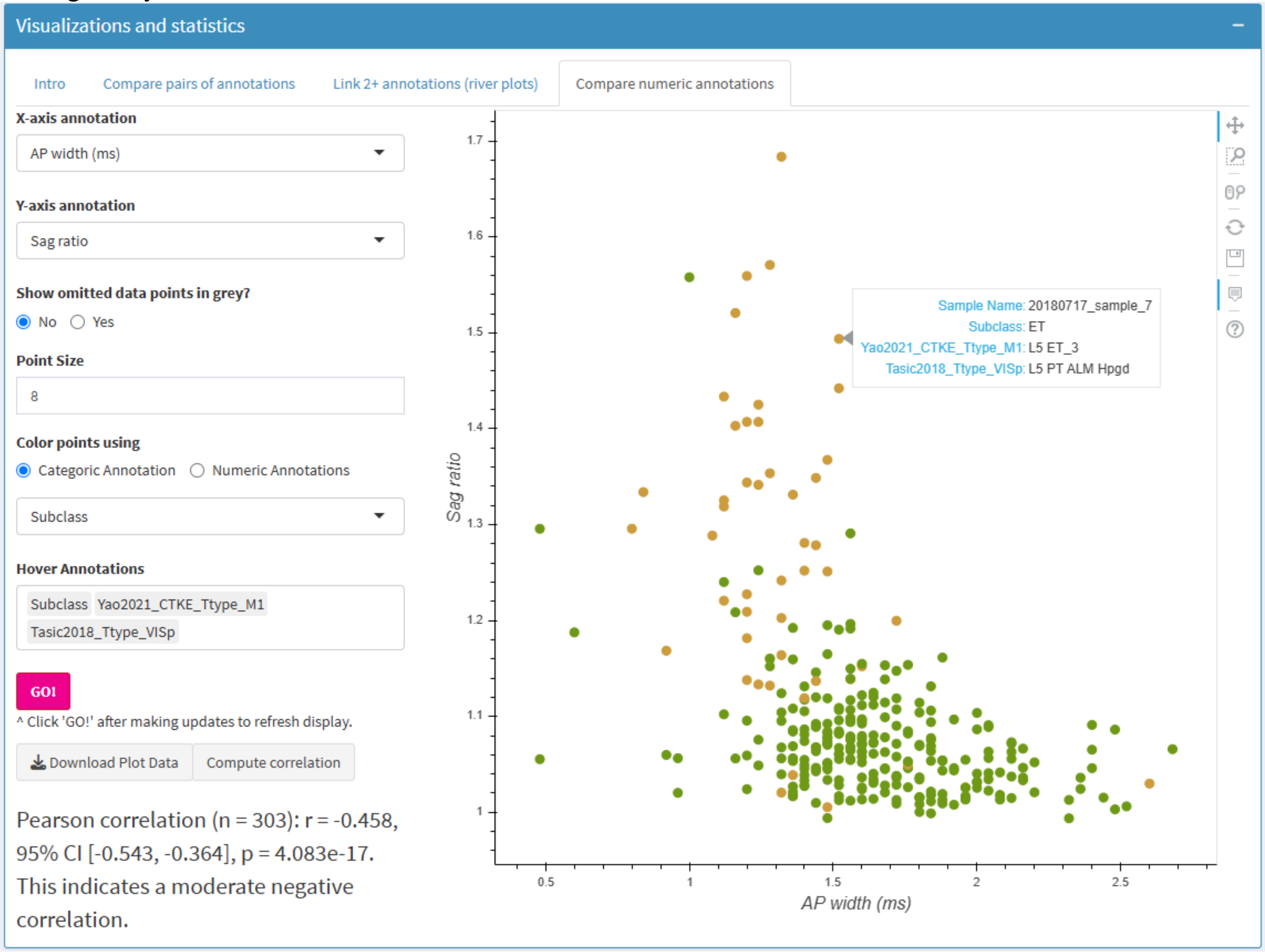


----------------------------

### Conclusions

Thank you for reading the user guide and for using ACE! Please reach out with any suggestions for how to improve the tool or associated documentation, or if you still have any questions or comments about the tool.

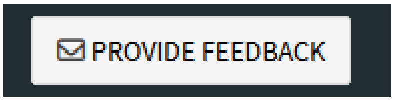


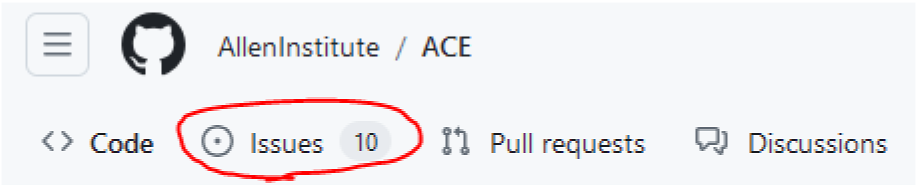
